# Breakage of Hydrophobic Contacts Limits the Rate of Passive Lipid Exchange Between Membranes

**DOI:** 10.1101/2020.05.06.081562

**Authors:** Julia R. Rogers, Phillip L. Geissler

**Affiliations:** Department of Chemistry, University of California, Berkeley, CA 94720, United States; Chemical Sciences Division, Lawrence Berkeley National Laboratory, Berkeley, CA 94720, United States

## Abstract

The maintenance of heterogeneous lipid compositions among cellular membranes is key to biological function. Yet, even the simplest process that could be responsible for maintaining proper lipid distributions, passive lipid exchange of individual molecules between membranes, has eluded a detailed understanding, due in part to inconsistencies between experimental findings and molecular simulations. We resolve these discrepancies by discovering the reaction coordinate for passive lipid exchange, which enables a complete biophysical characterization of the rate limiting step for lipid exchange. Our approach to identify the reaction coordinate capitalizes on our ability to harvest over 1,000 unbiased trajectories of lipid insertion, an elementary step of passive lipid transport, using all-atom and coarse-grained molecular dynamics simulations. We find that the reaction coordinate measures the formation and breakage of hydrophobic contacts between the membrane and exchanging lipid. Consistent with experiments, free energy profiles as a function of our reaction coordinate exhibit a substantial barrier for insertion. In contrast, lipid insertion was predicted to be a barrier-less process by previous computational studies, which incorrectly presumed the reaction coordinate to be the displacement of the exchanging lipid from the membrane. Utilizing our newfound knowledge of the reaction coordinate, we formulate an expression for the lipid exchange rate to enable a quantitative comparison with experiments. Overall, our results indicate that the breakage of hydrophobic contacts is rate limiting for passive lipid exchange and provide a foundation to understand the catalytic function of lipid transfer proteins.

## INTRODUCTION

Lipid membranes are essential to the structural integrity of cells. They form fluid boundaries that organize and compartmentalize cellular functions within organelles. Each organelle requires a unique membrane composition for its proper function. ^1,2^ To maintain such varied compositions, lipids are heterogeneously trafficked between cellular membranes through vesicular or non-vesicular transport. Vesicular transport plays a major role in trafficking lipids and proteins between organelles in the secretory pathway. ^3,4^ Organelles that are not connected by vesicular transport machinery rely on non-vesicular mechanisms to receive and export lipids. Even for organelles in the secretory pathway, non-vesicular transport mechanisms provide an additional way to more rapidly exchange lipids, for example, to swiftly alter membrane compositions in response to environmental changes. ^5,6^

Despite its cellular importance, non-vesicular transport mechanisms *in vivo* have not been fully characterized. Non-vesicular transport predominantly involves the movement of individual lipids between membranes. Monomeric lipid transfer between membranes may occur passively, in which a lipid desorbs and freely diffuses to another membrane. ^5^ Alternatively, lipid transfer proteins may facilitate monomeric lipid exchange by enclosing lipids within their hydrophobic interiors during transport. ^6,7^ With half-times on the order of hours, passive lipid exchange is too slow to fully account for lipid transport *in vivo*, ^5^ indicating that lipid transfer proteins are likely catalysts of lipid exchange. Nevertheless, a mechanistic understanding of passive lipid exchange can inform our knowledge of how lipid transfer proteins increase the rate of lipid transport.

Molecular simulations are well suited to explore the microscopic dynamics and to identify the rate limiting step of passive lipid exchange. Quantifying the free energy barrier associated with the rate limiting step, however, requires knowledge of the reaction coordinate, which characterizes the collective motion of molecules that advances a transition. ^8–11^ Previous computational work ^12–16^ on lipid transport has presumed that a lipid’s displacement normal to the membrane is the reaction coordinate (Figure 1A) and has yielded results in conflict with experimental findings. ^17–20^ Here we show that the reaction coordinate for passive lipid exchange is indeed more subtle than a simple distance measurement. The reaction coordinate characterizes the creation (or disruption) of a locally hydrophobic environment around the incoming (or outgoing) lipid (Figure 1B). This realization resolves qualitative (but not quantitative) discrepancies between simulation and experiment and suggests that the breakage of hydrophobic contacts between a lipid and membrane limits the rate of passive lipid transport.

**Figure 1:**
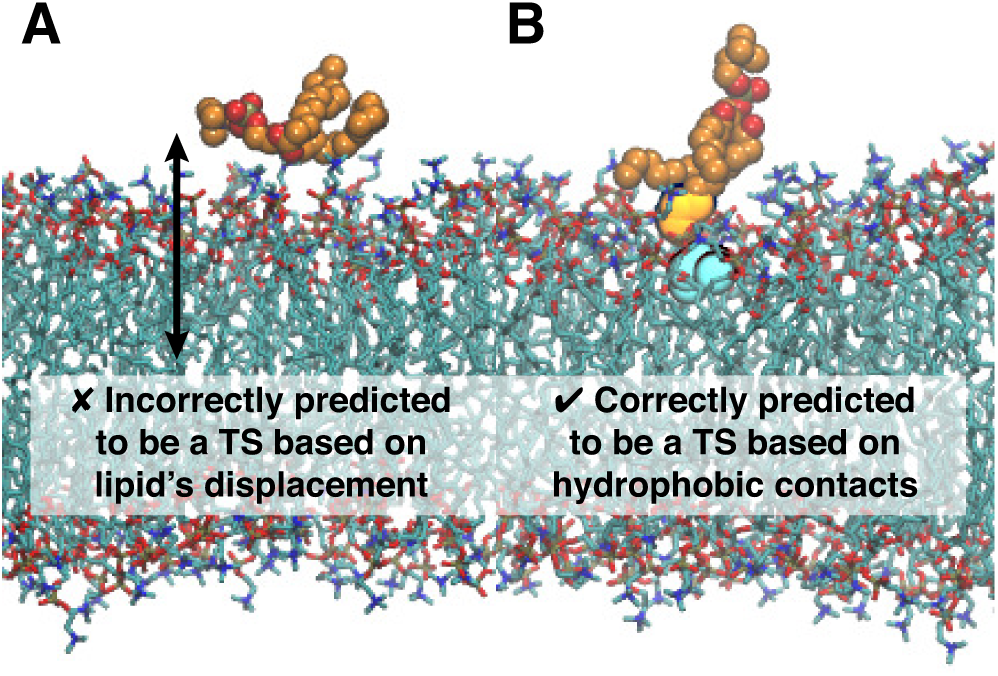
A lipid’s displacement normal to the membrane has conventionally been presumed to be the reaction coordinate for passive lipid exchange; however, the lipid’s displacement does not reliably identify transition state (TS) configurations. Although the lipid’s displacement (black arrow) in configuration A is consistent with values observed at the transition state, this configuration is not a transition state. By contrast, configuration B is correctly predicted to be a transition state based on the extent of hydrophobic contact between the lipid and membrane (highlighted in yellow and cyan).

### Experimental and Computational Background

Numerous *in vitro* studies of passive monomeric lipid exchange between membranes have demonstrated that it is a first-order process and that the rate of lipid exchange strongly correlates with a lipid’s solubility. ^17–19,21–27^ Based on these observations, the most widely accepted mechanism is characterized by aqueous diffusion. ^17–19,21,24–31^ The diffusive mechanism involves lipid desorption, which is rate limiting, followed by diffusion through solvent and insertion into another membrane. Based on experimentally measured activation energies, calculated activation free energies for lipid exchange exceed free energies for transferring a lipid from water to a membrane, indicating that there is a barrier for lipid desorption and insertion. ^17–20^ At this barrier, the desorbing lipid is hypothesized to only have the terminal carbons of its tails left within the membrane. ^17,28^ Thus, the activation free energy required to form the transition state has been attributed to the creation of a cavity in the membrane due to partial removal of a lipid and another cavity in the solvent to accommodate that lipid. ^17,32^

However, because the experimental methods currently used to study lipid exchange have coarse temporal and spatial resolution, molecular features of the transient transition state can only be hypothesized. Molecular dynamics (MD) simulations offer an attractive means to test these hypotheses since the necessary time and length scales are accessible. Additionally, a complete free energy profile can be obtained from MD simulations. By calculating the free energy as a function of the reaction coordinate, which describes the system’s dynamics during transitions between stable states, ^8–10^ free energies of activation can in principle be accurately quantified. If, however, the free energy is computed as a function of an order parameter that is not the reaction coordinate, then the apparent barrier generally underestimates the rate-determining free energy of activation. ^11^ Thus, identifying the reaction coordinate and corresponding free energy profile for passive lipid exchange can provide fundamental insights into the physical processes and work required to maintain heterogeneous cell membrane compositions.

Most previous computational studies have focused on obtaining a full free energy profile for lipid desorption and insertion. ^12–16^ Traditionally, free energy profiles have been computed as a function of a lipid’s displacement normal to a bilayer measured from the lipid’s phosphate group to the bilayer’s center-of-mass (COM) (Figure 1A). ^12–16^ These free energy profiles lack a barrier for insertion, ^12–16^ seemingly in conflict with experimental results. ^17–20^ If the COM displacement is not the reaction coordinate for lipid exchange, the kinetically relevant barrier may not be resolved in these free energy profiles; instead, a barrier may exist along a different degree of freedom that is the reaction coordinate. Consistent with this idea, Vermaas and Tajkhorshid’s MD study of lipid insertion indicated that the COM displacement is not sufficient to fully describe the microscopic dynamics of lipid insertion. They demonstrated that after the lipid associates with a bilayer, each tail of the lipid enters the bilayer successively to complete the insertion processes. The observation of splayed lipid intermediates during insertion, which are indistinguishable from other configurations based on the lipid’s COM displacement, suggests that other degrees of freedom need to be considered to construct an accurate reaction coordinate for lipid exchange. ^33^

### Our Approach

The discrepancy between experimental^17–20^ and computational ^12–16^ reports about a barrier for lipid exchange, together with the evidence suggesting that the lipid’s COM displacement is a poor reaction coordinate from a MD study of lipid insertion ^33^ prompt two questions: (1) What is the reaction coordinate for lipid exchange? (2) What, if any, activation free energy barrier impedes the process of lipid insertion? In this article, we aim to answer these questions using molecular simulation. In doing so, we identify the reaction coordinate for passive lipid exchange in all-atom and coarse-grained lipid models. Knowledge of the reaction coordinate allows us to elucidate key biophysical details of the transition state ensemble and properly assess the free energetic cost of lipid transport.

Rather than driving the system along a presumed reaction coordinate, we instead harvest natural, unbiased trajectories in which a lipid spontaneously inserts into a membrane. Statistical analysis of this ensemble of dynamical pathways reveals the key collective motions required for lipid transport. Firstly, we find that lipid insertion is a barrier crossing event and occurs *via* three different pathways, distinguished by various splayed lipid intermediates. Secondly, we find that the reaction coordinate characterizes the formation and breakage of hydrophobic lipid–membrane contacts. The lipid’s displacement normal to the bilayer, even formulated to distinguish splayed configurations, is not the reaction coordinate and obfuscates the barrier for insertion. Consistent with previous experimental results, ^17–20^ free energy profiles as a function of our reaction coordinate display a barrier for insertion and yield insertion rates calculated from Kramers theory that agree with those obtained directly from simulations. Finally, using our newfound reaction coordinate, we formulate a Smoluchoski equation for the lipid exchange rate to directly compare our simulation results with experiments. Overall, our results demonstrate that the rate limiting step for passive lipid exchange is the breakage of hydrophobic contacts, suggesting that lipid transfer proteins may catalyze lipid transport in part by lowering the associated activation free energy.

## METHODS

### Molecular Dynamics Simulations

Spontaneous desorption of a lipid from a membrane is a very slow process occurring over minutes to hours, which is well beyond the timescale accessible in MD simulations. Most previous work has addressed this problem by introducing an external bias that allows rare configurations otherwise inaccessible in MD simulations to be readily sampled. While this approach generates configurations plausibly found along a lipid desorption trajectory, it can fail to reveal the natural, unbiased route of lipid desorption. Our approach instead exploits a fundamental statistical property of microscopic dynamics, namely its time reversibility. Natural desorption trajectories are simply the time reverse of spontaneous lipid insertion trajectories. As shown by Vermaas and Tajkhorshid, ^33^ the latter are straightforward to generate in an unbiased way since lipid insertion is a rapid process. Thus, to gain insights into the dynamics of lipid exchange, we harvested trajectories of lipid insertion into a bilayer of 128 lipids using both all-atom and coarse-grained MD simulations. All simulations were performed in an isothermal-isobaric (NPT) ensemble using GROMACS 5. ^34^ The temperature was maintained at 320 K, ensuring that the bilayers were in the liquid crystalline phase. The pressure was maintained at 1 bar using semi-isotropic pressure coupling to allow the *z* dimension, which is perpendicular to the bilayer, to fluctuate separately of *x* and *y*, ensuring tensionless bilayers.

#### Coarse-Grained Systems

A large number of lipid insertion trajectories are required to make conclusions with high statistical accuracy. To obtain 1,000 lipid insertion trajectories in a computationally tractable manner, we performed MD simulations using the coarse-grained MARTINI force field. ^35^ All coarse-grained simulations used dilauroylphosphatidylcholine (DLPC) to compare to all-atom simulations of dimyristoylphosphatidylcholine (DMPC). Because MARTINI maps roughly four heavy atoms to a single coarse-grained bead, DMPC, which has 14 carbon atoms per tail, is best represented by the MARTINI model for DLPC, which has 3 beads per tail.

First, a bilayer surrounded by 3 nm thick slabs of standard MARTINI water was built using INSANE. ^36^ Prior to the addition of a tagged lipid into the solvent surrounding a bilayer of MARTINI DLPC lipids, the bilayer’s structure was fully relaxed from its initial lattice configuration. This involved an energy minimization using the steepest descent algorithm followed by equilibration and production runs. During the first 500 ps equilibration run, a 10 fs time step and the Berendsen barostat ^37^ with a coupling time constant of 3 ps and an isothermal compressibility of 3 × 10^−4^ bar^−1^ were used. During a second 1 ns equilibration run, the time step was increased to 30 fs and the barostat was switched to the Parinello-Rahman algorithm ^38^ with a coupling time constant of 12 ps. A 50 ns production run using the same simulation parameters as the second equilibration run was performed to allow the bilayer’s structure to fully equilibrate, as monitored by the area per lipid (Figure S1). The temperature was maintained at 320 K with the V-rescale thermostat^39^ using a coupling time constant of 1 ps. The lipids and solvent were coupled to separate thermostats to avoid the “hot solvent-cold solute” problem. ^40^ Dynamics were evolved according to the leapfrog algorithm. ^41^ As determined to yield optimal performance for simulations using MARTINI, ^42^ neighbor lists were updated using the Verlet neighbor searching algorithm, ^43^ Lennard-Jones and Coulomb interactions were truncated at 1.1 nm, and Coulomb interactions beyond the cutoff were evaluated with a reaction-field potential^44^ with a relative dielectric constant of *∞.*

Next, 1,000 replicate systems with a free lipid were built by inserting a tagged lipid at a random location in the solvent around the equilibrated bilayer such that the tagged lipid’s COM and bilayer’s COM are separated in *z* by at least 3.2 nm. All replicates were then energy minimized and equilibrated using the protocol described above with one modification: To ensure that the tagged lipid did not adsorb or insert into the bilayer during equilibration, the *z* coordinates of its heavy atoms were restrained by a harmonic potential with a force constant of 500 kJ/mol/nm^2^. Upon release of the position restraints, production runs of 1 *µ*s were performed, and lipid insertion occurred in all replicates during this time. Reported times for MARTINI simulations are not scaled by a factor of 4, as done in other work to account for the erroneously fast solvent diffusion in MARTINI as compared to real water. ^35^

#### All-Atom Systems

Additionally, we harvested 10 all-atom lipid insertion trajectories to compare with our results using MARTINI. The CHARMM36 force field ^45^ was used in combination with the CHARMM TIP3P water model ^46^ since it accurately reproduces many experimental observables, including the volume and area per lipid, bilayer thickness, lipid lateral diffusion coefficient, and neutron density profiles, for a liquid crystalline DMPC bilayer. ^47,48^ We used a simulation protocol similar to that described above for MARTINI.

First, a bilayer surrounded by 3 nm thick slabs of solvent was built using the CHARMM-GUI Membrane Builder. ^49,50^ The bilayer was energy minimized prior to undergoing a two-stage equilibration at 320 K and 1 bar. The first 250 ps equilibration utilized the Berendsen barostat^37^ for semi-isotropic pressure coupling with a coupling time constant of 2 ps and isothermal compressibility of 4.5 × 10^−5^ bar^−1^, and the second 250 ps equilibration utilized the Parinello-Rahman barostat ^38^ with a coupling time constant of 5 ps. A 50 ns production run was performed to allow the bilayer to fully equilibrate (Figure S1). The lipids and solvent were coupled to separate Nosé-Hoover thermostats^51,52^ using a coupling time constant of 1 ps to maintain the temperature. Dynamics were evolved according to the leapfrog algorithm ^41^ using a 2 fs time step. All bonds to hydrogen were constrained using the LINCS algorithm. ^53^ Lennard-Jones forces were smoothly switched off between 0.8 and 1.2 nm. Coulomb interactions were truncated at 1.2 nm, and long-ranged Coulomb interactions were calculated using Particle Mesh Ewald (PME) summation ^54^ with a Fourier spacing of 0.12 nm and an interpolation order of 4. Neighbor lists were constructed with the Verlet algorithm. ^43^

Next, 10 replicate systems with a tagged lipid in solution were built as for the MARTINI systems. Each replicate was energy minimized and equilibrated using the protocol described for the CHARMM36 bilayer system with the addition of harmonic restraints on the *z* coordinates of all heavy atoms of the tagged lipid. After the position restraints were removed, each replicate was simulated in increments of 100 ns until the tagged lipid inserted into the bilayer.

### Characterization of Transition Paths

From the harvested lipid insertion trajectories, we identified transition paths, trajectory segments that connect “reactant” and “product” states A and B. In state A, the tagged lipid is fully surrounded by solvent; in state B, the tagged lipid is fully within the bilayer. States A and B are characterized by the displacements *d*_lip_, *d*_sn1_, and *d*sn2 shown in Figure 2A. *d*lip is the displacement in *z* from the COM of the tagged lipid to the COM of the closest leaflet. Similarly, *d*_sn1_ is the displacement in *z* from the terminal carbon of the sn1 tail of the tagged lipid to the COM of the closest leaflet, and *d*_sn2_ is the analogous distance for the sn2 tail. Based on distributions of these three distances obtained from coarse-grained and all-atom MD simulations (Figure S2), state A is defined by *d*_lip_ > 24 Å, which ensures that any configurations with the tagged lipid adsorbed onto the surface of the bilayer are not included. State B is defined by *d*_sn1_ < −3 Å and *d*_sn2_ <− 3 Å, which ensures that the both tails of the tagged lipid are inserted into the bilayer.

**Figure 2:**
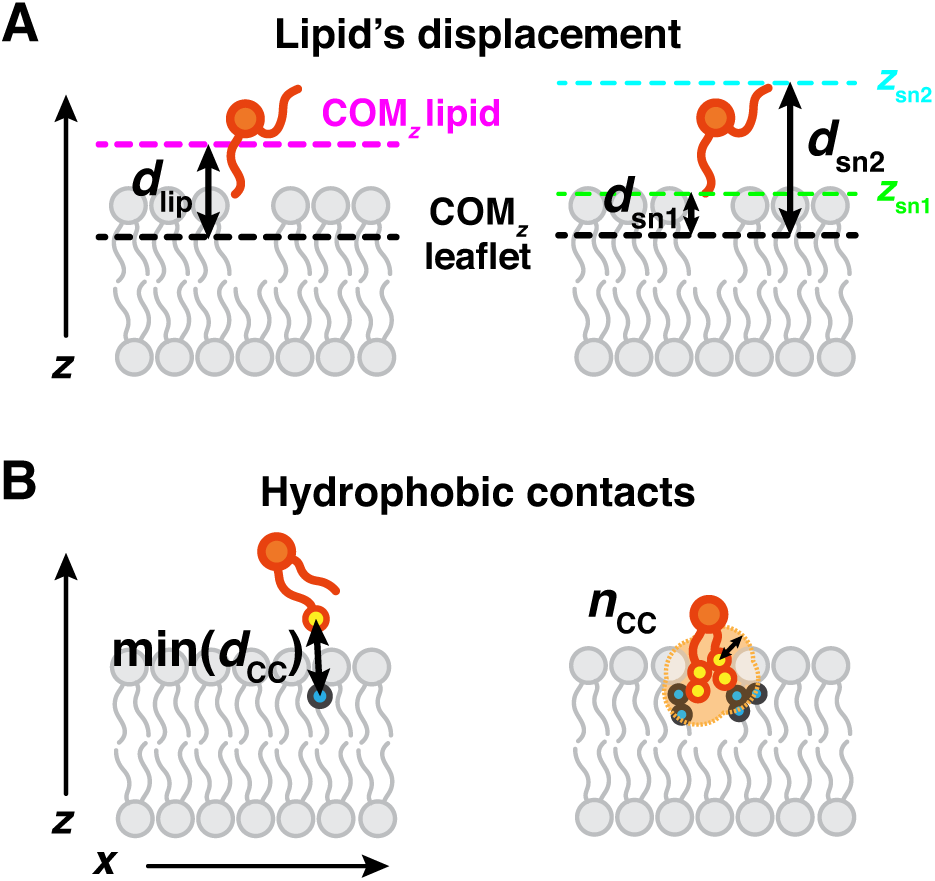
Schematics of order parameters that measure (A) the displacement of the lipid along the bilayer’s normal and (B) hydrophobic lipid–bilayer contacts. The tagged lipid is colored orange. (A) *d*_lip_ is the displacement in *z* between the center-of-mass (COM_*z*_) of the tagged lipid (magenta dashed line) and COM_*z*_ of the closest leaflet (black dashed line). *d*_sn1_ is the displacement in *z* between the terminal carbon of the sn1 tail (green dashed line) and COM_*z*_ of the closest leaflet. *d*_sn2_ is the displacement in *z* between the terminal carbon of the sn2 tail (cyan dashed line) and COM_*z*_ of the closest leaflet. (B) min(*d*_CC_) is the minimum distance between a hydrophobic carbon of the tagged lipid and a hydrophobic carbon of the closest leaflet. The closest pair of hydrophobic carbons are drawn as circles. *n*_CC_ is the total number of close hydrophobic carbon contacts between the tagged lipid and closest leaflet. Any pair of lipid and membrane hydrophobic carbons within a cutoff distance of 14 Å for MARTINI and 10 Å for CHARMM36 are counted as contacts and drawn as circles. The light orange region highlights the space within a cutoff distance (black arrow) from hydrophobic carbons of the tagged lipid.

In addition to *d*_lip_, *d*_sn1_, and *d*_sn2_, we evaluated over 50 order parameters as putative reaction coordinates for lipid exchange. Hydrophobic contacts between the tagged lipid and bilayer are judged according to the distance 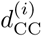 between a hydrophobic carbon of the tagged lipid and a hydrophobic carbon of the closest membrane leaflet, where *i* indexes the many pairs of such atoms. For MARTINI lipids, hydrophobic carbons include all tail beads. For CHARMM36 lipids, hydrophobic carbons include atoms C23 – C214 and C33 – C314. More specifically, hydrophobic contacts are measured by the order parameters min(*d*_CC_) and *n*_CC_ (Figure 2B). 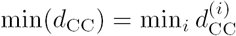 is the minimum distance between a hydrophobic carbon of the tagged lipid and a hydrophobic carbon of the closest leaflet. *n*_CC_ is the number of close contacts between hydrophobic carbons of the tagged lipid and hydrophobic carbons of the closest leaflet. The *i*th pair of hydrophobic carbons was counted as a close contact if 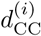 ≤ 14 Å for MARTINI lipids and if 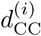 ≤ 10 Å for CHARMM36 lipids. These cutoff values encompass approximately two water solvation shells around a hydrophobic carbon of a lipid in solution and also two carbon solvation shells around a hydrophobic carbon of a lipid in a bilayer. We found smaller cutoffs to be insufficient for fully characterizing the tagged lipid’s hydrophobic environment and, thus, also insufficient for constructing the reaction coordinate. Our definition of *n*_CC_ is similar to the order parameter developed by Lin and Grossfield to measure hydrophobic contacts between lipopeptides and phospholipids. Similar to the conclusions we make herein about phospholipid exchange, they found that a hydrophobic contact order parameter was key to accurately investigate lipopeptide insertion from a micelle into a phospholipid bilayer, whereas the COM displacement was insufficient.^55^ Complete descriptions of all other order parameters are provided in the SI. The MDAnalysis Python library ^56^ was used to analyze all trajectories for each order parameter.

### Free Energy Calculations

Based on the dynamics observed along transition paths, lipid insertion is a barrier-crossing process. To identify the physical origin of this barrier, we calculated free energy surfaces as a function of different order parameters. To obtain these free energy surfaces, we performed umbrella sampling simulations ^57^ using the PLUMED 2 patch^58^ for GROMACS.

#### Coarse-Grained Systems

We computed the 2D free energy surfaces Δ*F* (min(*d*_CC_), *n*_CC_) and Δ*F* (*d*_sn1_, *d*_sn2_) for the MARTINI system. To obtain Δ*F* (min(*d*_CC_), *n*_CC_), we simulated 208 windows with harmonic biases centered at physically possible values of (min(*d*_CC_), *n*_CC_) (Table S1) for 2 *µ*s each. Each window was initialized with a configuration, drawn from a transition path, that has a value of (min(*d*_CC_), *n*_CC_) close to the center of the window’s bias. To calculate biasing forces on min(*d*_CC_), a smooth form for the minimum function

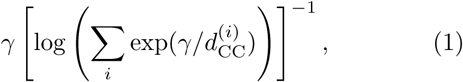

was used with *γ* = 260. min(*d*_CC_) calculated with expression 1 differed from the exact value by 0.007 Å on average and by at most 0.34 Å. To calculate biasing forces on *n*_CC_, a switching function was used

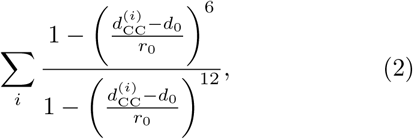

with *d*_0_ = 14 Å and *r*_0_ = 0.25 Å. *n*_CC_ calculated with expression 2 differed from the exact value by 4 contacts on average and by at most 28 contacts.

To obtain Δ*F* (*d*_sn1_, *d*_sn2_), we simulated 370 windows with harmonic biases centered at positions (*d*_sn1_, *d*_sn2_) ranging from (−10 Å, −10 Å) to (30 Å, 30 Å) for 2 *µ*s each. To avoid unrealistically distorting the tagged lipid, we simulated only windows whose harmonic bias centers satisfy |*d*_sn1_ − *d*_sn2_ | ≤30 Å. Each window was initialized with a configuration, drawn from a transition path, that has a value of (*d*_sn1_, *d*_sn2_) close to the center of the window’s bias. A force constant of 500 kJ/mol/nm^2^ was used for all harmonic bias potentials.

The weighted histogram analysis method (WHAM) ^59^ was used to obtain both of these 2D free energy surfaces from the biased distributions, after discarding data from the first 1 *µ*s. Error bars were calculated as the standard error of free energy surfaces estimated from five independent 200 ns blocks.

#### All-Atom Systems

To obtain Δ*F* (min(*d*_CC_), *n*_CC_) for the CHARMM36 system, 202 windows with harmonic biases centered at physically possible values of (min(*d*_CC_), *n*_CC_) (Table S2) were simulated for 24 ns each. Expression 1 was used to calculate biasing forces on min(*d*_CC_) with *γ* = 200, and it differed from the exact value by 0.002 Å on average and by at most 0.27 Å. Expression 2 was used to calculate biasing forces on *n*_CC_ with *d*_0_ = 10 Å and *r*_0_ = 0.25 Å, and it differed from the exact value by 68 contacts on average and by at most 175 contacts. Each window was initialized with a configuration, drawn from a transition path, that has a value of (min(*d*_CC_), *n*_CC_) close to the center of the window’s bias. The first 4 ns of data from these windows was discarded to account for equilibration. Finally, data from all windows was combined with WHAM to obtain a free energy surface as a function of min(*d*_CC_) and *n*_CC_. Error bars were calculated as the standard error of free energy surfaces estimated from five independent 5 ns blocks.

#### Calculation of 1D Free Energy Profiles from 2D Free Energy Surfaces

We calculated 1D free energy profiles Δ*F* (min(*d*_CC_)) and Δ*F* (*n*_CC_) by numerically integrating Δ*F* (min(*d*_CC_), *n*_CC_) over one of its variables. Denoting the two variables as *q* and *q*′ (in either order), the free energy profile Δ*F* (*q*) is obtained from the free energy surface Δ*F* (*q, q*′) according to

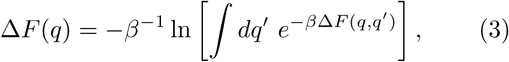

where *β* = (*k*_B_*T*)^−1^ is the inverse of Boltzmann’s constant, *k*_B_, multiplied by temperature.

### Committor Analysis

To identify the reaction coordinate for passive lipid exchange and determine if transition states are sampled in umbrella sampling simulations, we characterized configurations according to their tendency to proceed to state B using committor analysis. ^8–10^ The committor, *p*_B_, is the probability that a configuration will reach state B prior to state A when its momenta are chosen randomly from a Maxwell-Boltzmann distribution. By construction, the committor distinguishes transition states, which have *p*_B_ = 0.5, from stable state A and B configurations, which have *p*_B_ = 0 and 1, respectively. Thus, *p*_B_ is the true reaction coordinate. Through committor analysis of configurations found along transition paths, we identified order parameters that are strongly correlated with the committor and, thus, can be used as approximate reaction coordinates. These order parameters have the advantage of being more physically descriptive and, thus, more easily interpreted than the committor. Henceforth we refer to order parameters that correlate strongly with *p*_B_ as the reaction coordinate. We calculated committor values for 98,094 MARTINI and 138 CHARMM36 configurations sampled along transition paths. Additionally, we calculated the committor for 500 MARTINI and 100 CHARMM36 configurations from each umbrella sampling simulation. For each MARTINI configuration, the outcome of 50 trajectories, each 3 ns long and initialized with random velocities sampled from a Maxwell-Boltzmann distribution, were used to calculate its committor value. For each CHARMM36 configuration, the outcome of 20 trajectories, each 12 ns long, were used to calculate its committor value.

### Rate Calculations

To further assess how well the dynamics of lipid transport are captured by monitoring lipid–membrane hydrophobic contacts, we calculated the rate constant for lipid insertion, *k*_ins_, using Kramers theory^60^ coupled with thermodynamic information about hydrophobic contacts. The resulting value was compared with the insertion rate calculated from the mean first passage time in MD simulations. Based on Kramers theory,

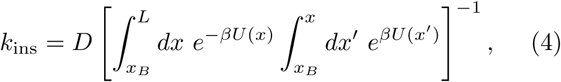

where *D* is the diffusion coefficient along a reaction coordinate *x* for lipid insertion, *x*_*B*_ is the value of *x* in state B, *L* is the maximal value of *x* in state A sampled in our simulations, and *U* (*x*) is the effective interaction potential biasing the dynamics of *x*. We have taken *x* to be min(*d*_CC_) since min(*d*_CC_) describes the spatial motion of the lipid during insertion in addition to hydrophobic contact formation. For the same reason we set *D* to be the diffusion coefficient of a freely diffusing lipid in solution. Note that min(*d*_CC_) alone is not the reaction coordinate for lipid exchange (a linear combination of min(*d*_CC_) and *n*_CC_ is the reaction coordinate), but it is more simply interpretable to utilize a single order parameter for these calculations. *U* (*x*) is taken to be the free energy profile Δ*F* (min(*d*_CC_)). *U* (*x*) was obtained from the free energy surface that depends jointly on min(*d*_CC_) and *n*_CC_ using Eq. 3. We evaluated Eq. 4 with numerical integration using the trapezoidal rule. The diffusion coefficient was calculated from an additional MD simulation of a single lipid solvated in a cubic box with the same area and simulation protocol as the bilayer systems, but with isotropic pressure coupling. *D* was calculated according to Einstein’s relation from the mean squared displacement of the lipid’s COM obtained from 1 *µ*s and 100 ns trajectories of a MARTINI and CHARMM36 lipid, respectively.

## RESULTS AND DISCUSSION

### Lipid Insertion Is a Barrier Crossing Process that Occurs *via* Multiple Pathways

We first investigated the dynamics of lipid insertion by harvesting 1,000 MARTINI DLPC and 10 CHARMM36 DMPC insertion trajectories from MD simulations. A lipid insertion event is classified by a transition from state A, in which the tagged lipid is fully in the solvent, to state B, in which the tagged lipid resides within the bilayer. State A is distinguished by *d*_lip_, which measures the COM displacement of the tagged lipid along the bilayer’s normal; state B is distinguished by *d*_sn1_ and *d*_sn2_, which measure the displacement of each tail of the tagged lipid along the bilayer’s normal (Figure 2A). Precise definitions are given above in the **Methods** section. The fact that state B configurations cannot be reliably identified by *d*_lip_ alone suggests that the COM displacement is not the reaction coordinate for passive lipid exchange.

Free energy profiles as a function of the lipid’s displacement along the bilayer normal obtained from previous computational studies ^12–16^ give the impression that lipid insertion is a barrier-less process. If that were true, insertion should occur immediately once the tagged lipid reaches the bilayer. However, as seen in snapshots from a MARTINI and a CHARMM36 trajectory shown in Figure 3 and Movies S1 and S2, the tagged lipid repeatedly arrives at the bilayer and adheres to its surface without inserting. Instead, it detaches from the bilayer’s surface and returns to the solvent. In typical trajectories, many such unproductive encounters occur before the lipid inserts into the bilayer. Indeed, as seen in time traces of *d*_lip_ in Figure 4A, many adsorption events commonly precede insertion. Similar adsorption events have been observed during simulations of 1-palmitoyl-2-oleoylphosphatidylcholine (POPC) insertion into a bilayer. ^33^

**Figure 3:**
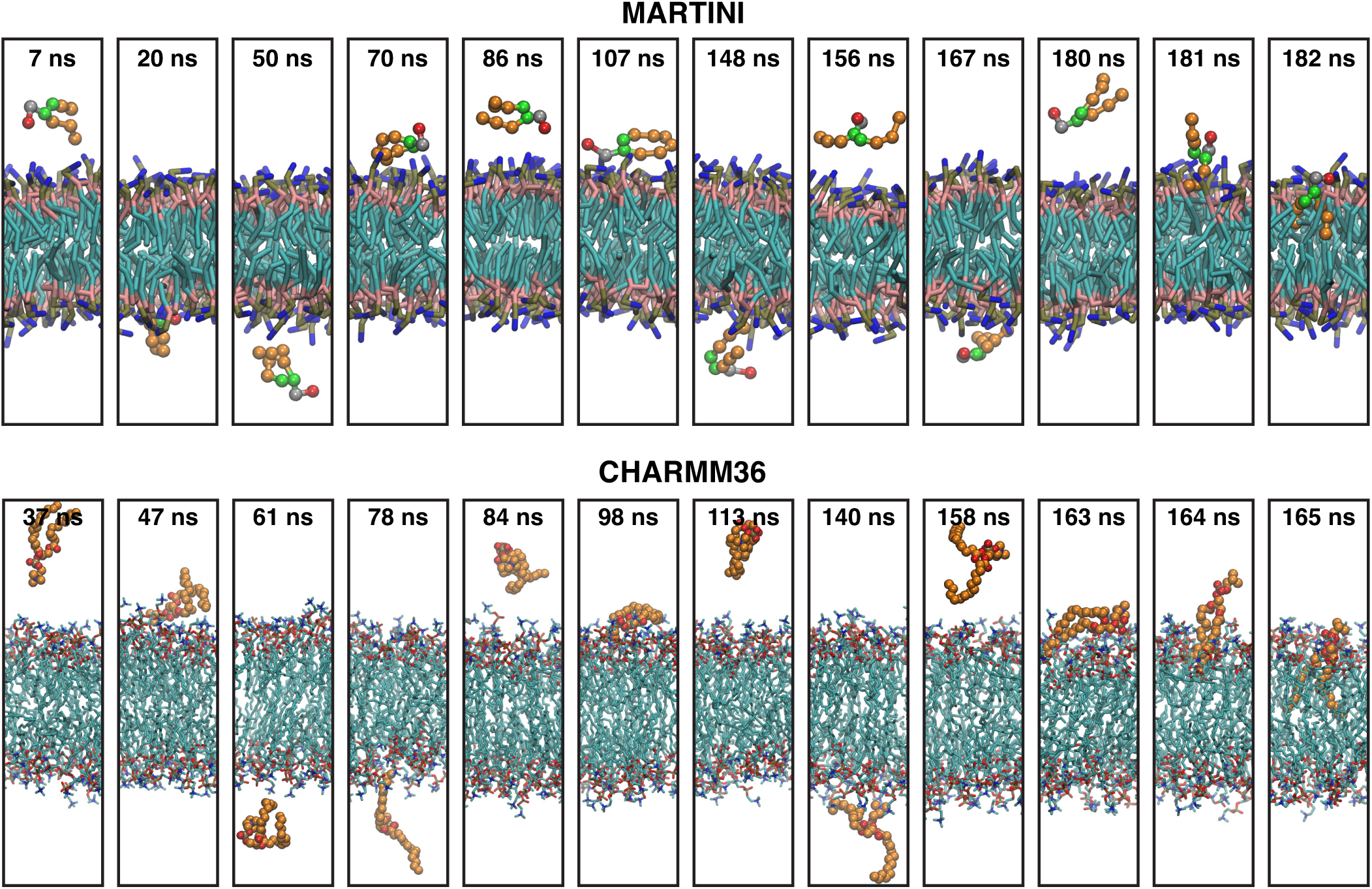
Snapshots along MD trajectories of a tagged lipid inserting into a bilayer illustrate that a membrane-adsorbed lipid does not immediately insert into a bilayer, suggesting that there is a barrier for insertion. For clarity, solvent is not shown in the snapshots. The tagged lipid is rendered with van der Waals spheres. For the MARTINI simulation, the headgroup, phosphate, glycerol, and tail beads of the tagged lipid (bilayer lipids) are colored red (blue), gray (brown), green (pink), and orange (cyan), respectively. For the CHARMM36 simulation, the carbon atoms of the tagged lipid (bilayer lipids) are colored orange (cyan).

**Figure 4:**
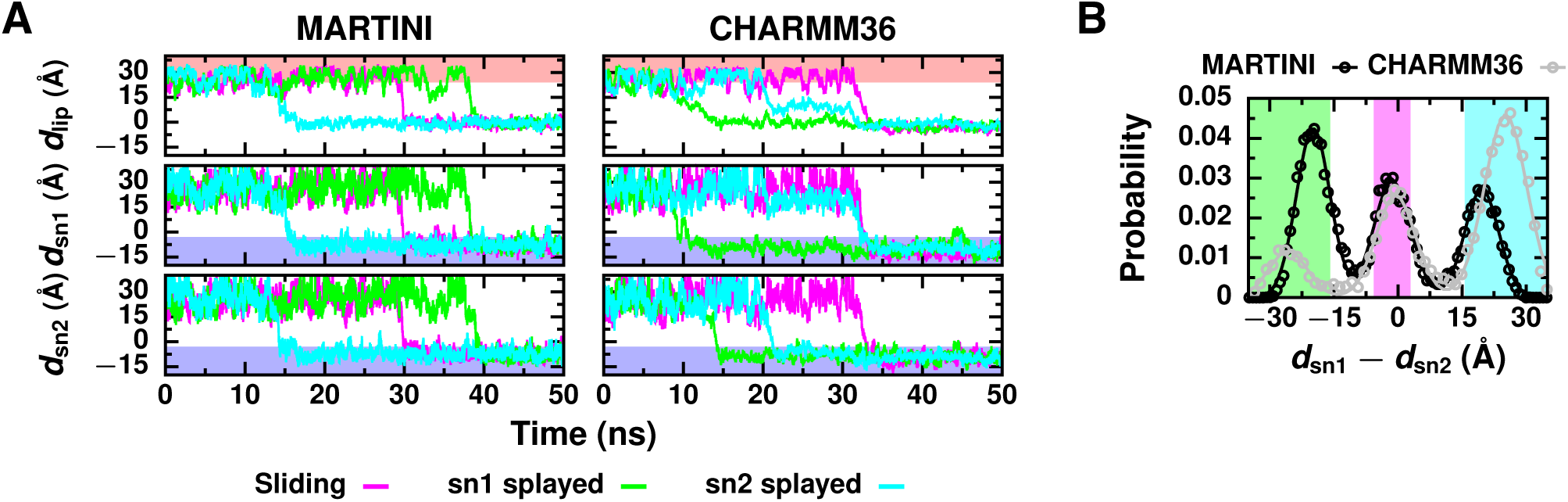
(A) Time evolution of *d*_lip_, *d*_sn1_, and *d*_sn2_ during MD simulations of lipid insertion is indicative of barrier crossing dynamics. State A configurations (*d*_lip_ > 24 Å) are located in the red region. State B configurations (*d*_sn1_ < −3 Å and *d*_sn2_ < −3 Å) are located in the blue region. Dips in the distances *d*_lip_, *d*_sn1_, and *d*_sn2_ out of the state A region without entering the state B region occur when the tagged lipid adsorbs to the surface of the bilayer. Examples for the sliding, sn1 splayed, and sn2 splayed pathways are shown for both MARTINI and CHARMM36. The initial times have been shifted so that all trajectories can be plotted together. (B) Probability distributions of *d*_sn1_ − *d*_sn2_ when the tagged lipid is near the surface of the bilayer (specifically, MARTINI configurations with 7 ≤ *d*_lip_ ≤ 9 Å and CHARMM36 configurations with 2.5 ≤ *d*_lip_ ≤ 15 Å), which were used to identify splayed tail configurations. Distributions from MD simulations are plotted with open circles, and the fits to a sum of three Gaussians are plotted with solid lines. Transition paths that follow the sn1 splayed pathway sample configurations found in the green region, and those that follow the sn2 splayed pathway sample configurations found in the cyan region. Transition paths that follow the sliding pathway sample configurations in the magenta region and do not sample splayed configurations in either the cyan or green regions.

When lipid insertion eventually occurs in these trajectories, it does so suddenly. Characteristic of barrier-crossing dynamics, *d*_lip_, *d*_sn1_, and *d*_sn2_ change sharply from values of state A to those of state B (Figure 4A). The fact that the transition times are much faster than the inverse rate constant for lipid insertion, 1*/k*_ins_, points to a substantial free energy barrier for insertion. Free energy profiles as a function of the lipid’s displacement simply do not resolve this barrier. ^12–16^ Thus, the barrier must exist along a different degree of freedom that captures other important features of the dynamics.

In fact, lipid insertion occurs *via* three different pathways which cannot be differentiated by *d*_lip_. Each pathway is characterized by a distinct lipid configuration, which is distinguished by *d*_sn1_ and *d*_sn2_, near the bilayer’s surface: (1) In the sliding pathway, the two tails enter the bilayer almost simultaneously as the tagged lipid slides into the bilayer (Figure 3, CHARMM36 trajectory and Figure 4A, magenta time traces). Near the bilayer’s surface, both tails are a similar distance from the bilayer (Figure 4B, magenta region). (2) In the sn1 splayed pathway, the sn1 tail enters the bilayer first (Figure 4A, green time traces), creating a splayed intermediate with the sn1 tail anchored in the bilayer (Figure 4B, green region). (3) In the sn2 splayed pathway, the sn2 tail enters the bilayer first (Figure 3, MARTINI trajectory and Figure 4A, cyan time traces), creating a splayed intermediate with the sn2 tail anchored in the bilayer (Figure 4B, blue region). Table S3 reports the frequency of each pathway in our simulations. Splayed intermediates have also been observed in a previous study of lipid insertion ^33^ and postulated as transition states for stalk formation during membrane fusion, ^61–66^ a key step in vesicular transport. The existence of distinct insertion pathways might suggest that the displacements of individual tails along the bilayer’s normal could serve as reaction coordinates since they encapsulate dynamically relevant information that is not contained in the COM displacement. We demonstrate below, however, that the displacements of individual tails are not the reaction coordinates for lipid exchange.

### The Reaction Coordinate Characterizes Hydrophobic Contacts Between the Lipid and Membrane

Based on experiments, a cavity model has been proposed to describe the transition state. ^17,32^ According to the cavity model, solvent is evacuated above the desorbing lipid and a void forms in the membrane below. Such a focus on cavities is reminiscent of modern theories of the hydrophobic effect, which characterize hydrophobicity in terms of the statistics of solvent density fluctuations. ^67^ Although no true cavities are observed at transition states sampled in our simulations (Figure S3), the importance of hydrophobicity in lipid transport is evident from our analysis of over 50 order parameters. For example, the number of water molecules solvating the tagged lipid steadily decreases during insertion (Figure S4). The density of hydrophobic molecular fragments below the tagged lipid gradually increases while the number of hydrophilic ones decreases (Figure S3). Defects in the polar head group region of the bilayer that expose hydrophobic membrane patches ^68,69^ are also observed in the transition state ensemble (Figures S5 and S6). Based on these results, we hypothesized that the reaction coordinate for passive lipid exchange monitors the formation and breakage of hydrophobic contacts between the tagged lipid and membrane (Figure 1B).

To rigorously test this hypothesis, we employed committor analysis. ^8–10^ The committor, *p*_B_, is the probability that a trajectory initiated at a given configuration will reach state B prior to state A when initial momenta are chosen randomly from a Maxwell-Boltzmann distribution. Configurations within stable states A and B have *p*_B_ = 0 and 1, respectively. Transition states are equally likely to advance to states B and A, such that *p*_B_ = 0.5. ^8–10,70^ Since *p*_B_ directly measures the progress of a reaction, *p*_B_ is the true reaction coordinate. ^9,10,71^ However, *p*_B_ is a complicated function of the system’s microscopic configuration — a very large set of variables in the case of biomolecular systems. The complete functional form for *p*_B_ is practically unobtainable and would provide little physical insight into reaction mechanisms. Instead, it is more informative to identify an order parameter, *q*, that is strongly correlated with *p*_B_ and closely approximates the true reaction coordinate. ^71,72^ Henceforth we refer to such order parameters as the reaction coordinate.

As one assessment of correlation between an order parameter *q* and the true reaction coordinate *p*_B_, we compare the probability distribution of *q* from transition states to distributions of *q* from states A and B. The typical values of *q* at the transition state should differ from its values in states A and B such that these probability distributions do not overlap. Otherwise, *q* cannot reliably distinguish transition states from stable reactant and product states and, thus, poorly recapitulates the true reaction coordinate. Additionally, we compare the probability distribution of *q* from transition states to distributions from pre- and post-transition states. Because pre-transition states are intermediates between state A and the transition state and post-transition states are intermediates between the transition state and state B, it is often challenging to devise an order parameter *q* that distinguishes transition states from them, as *p*_B_ naturally does. A good approximation to the true reaction coordinate should reliably distinguish transition states from pre- and post-transition states in addition to state A and B configurations.

A more stringent test of a putative reaction coordinate examines a histogram of *p*_B_ values for configurations with a particular value of *q*. If *q* is the reaction coordinate, then a histogram of *p*_B_ for configurations with a given value of *q* will be peaked at the corresponding value of *p*_B_. This histogram test is useful to determine if the mapping from *q* to *p*_B_ is approximately one-to-one, a requirement for *q* to be the reaction coordinate. Bimodal histograms of *p*_B_ are clear indicators that *q* is not the reaction coordinate. By contrast, a histogram sharply peaked at *p*_B_ = 0.5 for values of *q* characteristic of transition states clearly indicates that *q* accurately describes the true reaction coordinate. ^9,10^

Using the first criterion that *q* must reliably distinguish transition states from all other configurations to be the reaction coordinate, we assessed measures of hydrophobic lipid–membrane contacts as approximations to the true reaction coordinate. Specifically, we characterize hydrophobic contacts by the minimum distance between hydrophobic carbons of the tagged lipid and hydrophobic carbons of the bilayer, min(*d*_CC_), and the number of close hydrophobic carbon–carbon contacts between the tagged lipid and bilayer, *n*_CC_ (Figure 2B). Precise definitions are given above in the **Methods** section. To determine if this criterion was satisfied, we compared the probability distributions of min(*d*_CC_) and *n*_CC_ from several ensembles: equilibrium configurations representative of (1) state A and (2) state B (as defined above in **Methods**); and three ensembles drawn from transition paths of (3) pre-transition state configurations identified by *p*_B_ = 0, (4) transition states identified by *p*_B_ ≈ 0.5 (specifically, 0.45 ≤ *p*_B_ ≤ 0.55 for MARTINI and 0.4 ≤ *p*_B_ ≤ 0.6 for CHARMM36 configurations), and (5) post-transition state configurations identified by *p*_B_ = 1. Joint distributions of min(*d*_CC_) and *n*_CC_ in these five different ensembles are shown in Figure 5. Corresponding 1D probability distributions of min(*d*_CC_) and *n*_CC_ are shown in Figures S7 and S8. The distributions from MARTINI and CHARMM36 configurations exhibit similar features. In state A, min(*d*_CC_) effectively measures the separation between the tagged lipid and the distant bilayer. Large values of min(*d*_CC_) are therefore typical. No close hydrophobic contacts are formed since the tagged lipid is fully solvated. As the tagged lipid progresses from state A towards the transition state, it becomes a pre-transistion state configuration. Pre-transition state configurations are closer to the bilayer than state A configurations, resulting in decreased values of min(*d*_CC_), but hydrophobic contacts between the tagged lipid and bilayer still scarcely exist. At the transition state, the tagged lipid has made only a few initial hydrophobic contacts with the bilayer. The transition state distribution is centered at values of min(*d*_CC_) and *n*_CC_ intermediate between those of states A and B. As the tagged lipid progress from the transition state to state B, it becomes a post-transition state configuration. Post-transition state configurations, which include splayed lipid configurations, have a substantial number of hydrophobic contacts between the tagged lipid and bilayer compared to transition states but on average half as many as state B configurations. Both the post-transition state and state B distributions of min(*d*_CC_) are sharply peaked at a value consistent with the minimum of the Lennard-Jones potential for a carbon–carbon interaction (or a bead–bead interaction in the MARTINI model). In state B, a maximal number of hydrophobic contacts exist between the tagged lipid and adjacent lipids in the bilayer. Importantly, the transition state distribution overlaps negligibly with the other four distributions. A combination of min(*d*_CC_) and *n*_CC_ distinguishes transition states from not only stable state A and B configurations but also pre- and post-transition state configurations, and, therefore, may serve as the reaction coordinate for lipid exchange.

**Figure 5:**
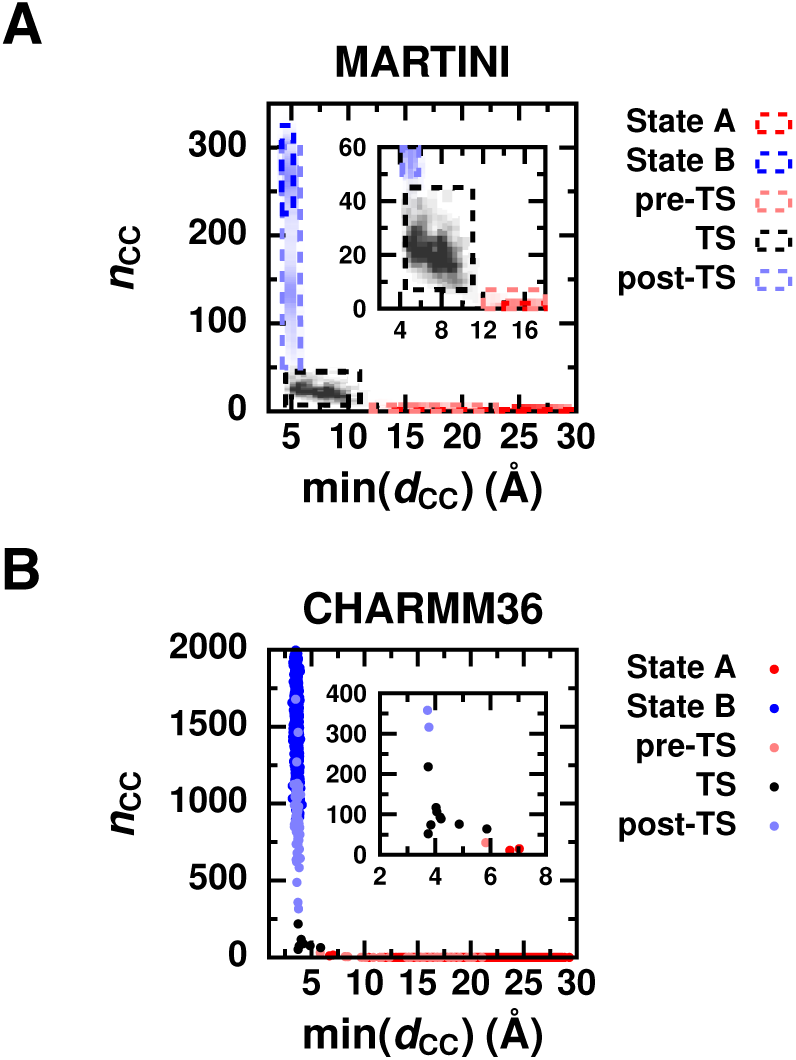
Distributions of min(*d*_CC_) and *n*_CC_ demonstrate that hydrophobic lipid–membrane contacts reliably identify transition states. (A) 2D probability distribution of min(*d*_CC_) and *n*_CC_ from MARTINI simulations. Individual 2D distributions for state A, for state B, and for three ensembles of configurations drawn from transition paths: pre-transition state (pre-TS) configurations, transition states (TS), and post-transition state (post-TS) configurations are each plotted in a single color and outlined in a dashed rectangle. (B) Scatter plot of min(*d*_CC_) and *n*_CC_ from CHARMM36 simulations. Magnified views around transition states are shown in the insets.

If the true reaction coordinate is well described by a combination of min(*d*_CC_) and *n*_CC_ — in other words if the formation and breakage of hydrophobic contacts are the essential processes required for lipid exchange — the free energy surface Δ*F* (min(*d*_CC_), *n*_CC_) should exhibit a barrier for insertion and desorption. These free energy surfaces for both MARTINI and CHARMM36 are shown in Figure 6A. Corresponding 1D free energy profiles along min(*d*_*CC*_) and *n*_*CC*_ are shown in Figure S10. A deep free energy minimum exists at values of min(*d*_*CC*_) and *n*_*CC*_ characteristic of state B due to the formation of many favorable hydrophobic contacts between the tagged lipid and membrane lipids. At larger values of min(*d*_CC_) characteristic of state A, the free energy surface flattens; when the lipid is far away from the membrane, the free energy is no longer sensitive to min(*d*_CC_). The surface plateaus at larger free energy values for MARTINI compared to CHARMM36. This discrepancy is consistent with the fact that compared to atomistic models, the MARTINI model overestimates the free energy difference between a lipid in solution and in the bilayer. ^35^ In qualitative agreement with experimental findings, ^17–20^ there is a free energy barrier for lipid insertion (Figure 6A, outlined in dashes). The activation free energy for the formation of hydrophobic contacts is roughly 5 *k*_B_*T*. Additionally, transition paths closely follow the minimum free energy path in the space of min(*d*_CC_) and *n*_CC_ (Figure S9), indicating that the dynamics of lipid exchange are well described in terms of hydrophobic contacts.

**Figure 6:**
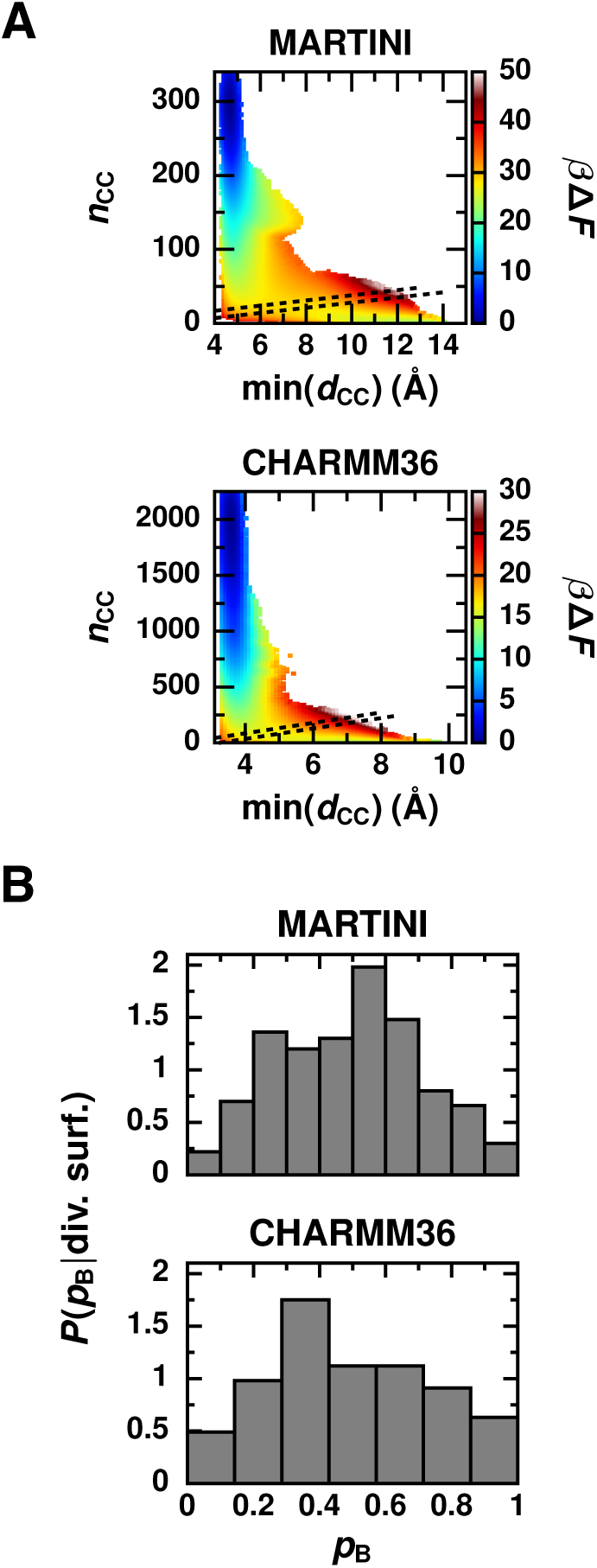
(A) Free energy surfaces as a function of min(*d*_CC_) and *n*_CC_ for the MARTINI and CHARMM36 force fields exhibit a barrier for insertion, which separates states A (bottom right corner) and B (top left corner). In each free energy profile, the dashed lines outline a narrow range about the dividing surface between states A and B. Specifically, the dashed lines outline the region −0.2 ≤ *r*_*c*_ ≤ 0.2 for MARTINI and −0.12 ≤ *r*_*c*_ ≤ 0.12 for CHARMM36, where *r*_*c*_ = *α*_1_ min(*d*_CC_) + *α*_2_*n*_CC_ + *α*_0_ with coefficients *α*_*i*_ determined using a maximum likelihood approach ^73,74^ (Table S5). Statistical errors in the free energy profiles are shown in Figure S9. (B) Histogram of committor values, *p*_B_, for configurations drawn from umbrella sampling simulations within a narrow range about the dividing surface, demonstrate that a linear combination of min(*d*_CC_) and *n*_CC_ is indeed the reaction coordinate.

Most importantly, a combination of min(*d*_CC_) and *n*_CC_ can locate transition states with high fidelity. We find that a linear combination suffices for this purpose (*r*_*c*_ = *α*_1_ min(*d*_CC_) + *α*_2_*n*_CC_ + *α*_0_ with coefficients *α*_*i*_ determined using a maximum likelihood approach ^73,74^ as detailed in the SI). By construction, *r*_*c*_ = 0 at the dividing surface between states A and B where transition states are located. The dashed lines in Figure 6A outline a narrow range around *r*_*c*_ = 0, which roughly traces the ridgeline of Δ*F* (min(*d*_CC_), *n*_CC_) between states A and B. To definitively test if *r*_*c*_ is the reaction coordinate, we performed committor analysis of configurations drawn from a narrow range around *r*_*c*_ = 0. Figure 6B shows that the ensemble defined by *r*_*c*_ ≈ 0 predominantly includes transition states for both MARTINI and CHARMM36. Thus, a measure of hydrophobic contacts between the tagged lipid and bilayer is the reaction coordinate for lipid exchange.

Together, min(*d*_CC_) and *n*_CC_ capture many different aspects of the lipid’s hydrophobic environment, underlying the ability of *r*_*c*_ to precisely describe the process of lipid exchange. Hydrophobicity can be quantified in other ways as well. For example, we constructed a more complicated reaction coordinate, 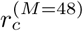, that is a linear combination of 48 order parameters excluding min(*d*_CC_) and *n*_CC_, with coefficients again determined using a maximum likelihood approach ^73,74^ (Table S4). As fully described in the SI, each of these 48 order parameters measure different details about the lipid’s environment, including the number of water molecules solvating the tagged lipid and the size of exposed hydrophobic membrane defects near the tagged lipid. This more complex reaction coordinate identifies transition states almost as faithfully as *r*_*c*_ does (Figure S11A), but 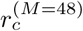 is difficult to interpret physically. The reaction coordinates *r*_*c*_ and 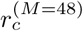 are highly correlated (Figure S11B), demonstrating that detailed information about the lipid’s environment is well represented by a simple combination of min(*d*_CC_) and *n*_CC_. Not only is a linear combination of min(*d*_CC_) and *n*_CC_ the most accurate reaction coordinate out of all tested (Table S5), but it has the advantage of providing physical insight into lipid exchange: The rate limiting step for desorption is the breakage of hydrophobic contacts between the lipid and membrane.

### The Lipid’s Displacement Normal to the Bilayer Is Not the Reaction Coordinate

Previous computational studies presumed that the lipid’s displacement normal to the bilayer is the reaction coordinate for lipid desorption and insertion. ^12–16^ However, *d*_lip_ is not the reaction coordinate, a fact we confirm by performing committor analysis. The probability distributions of *d*_lip_ from transition states are compared to the distributions from state A, state B, pre-transition state, and post-transition state configurations in Figure 7. The distributions from MARTINI and CHARMM36 configurations are quite similar. In state A, where the tagged lipid is fully solvated and away from the bilayer, the typical value of *d*_lip_ is large. As the lipid enters the bilayer, it progresses from being in state A to being a pre-transition state configuration, a transition state, a post-transition state, and finally in state B; correspondingly, the centers of each distribution shift to smaller values of *d*_lip_. In state B, the tagged lipid is in register with the other lipids in the membrane such that the distribution is centered near zero. For both MARTINI and CHARMM36, distributions from transition states overlap significantly with those from the other ensembles, indicating that transition states cannot be reliably identified by *d*_lip_ (Figure 1A). Furthermore, histograms of *p*_B_ for configurations sampled along transition paths with values of *d*_lip_ typical of transition states are bimodal and lack a peak at 0.5 (Figure S12), indicating that *d*_lip_ is not strongly correlated with *p*_B_. Therefore, *d*_lip_ is not the reaction coordinate for lipid exchange.

**Figure 7:**
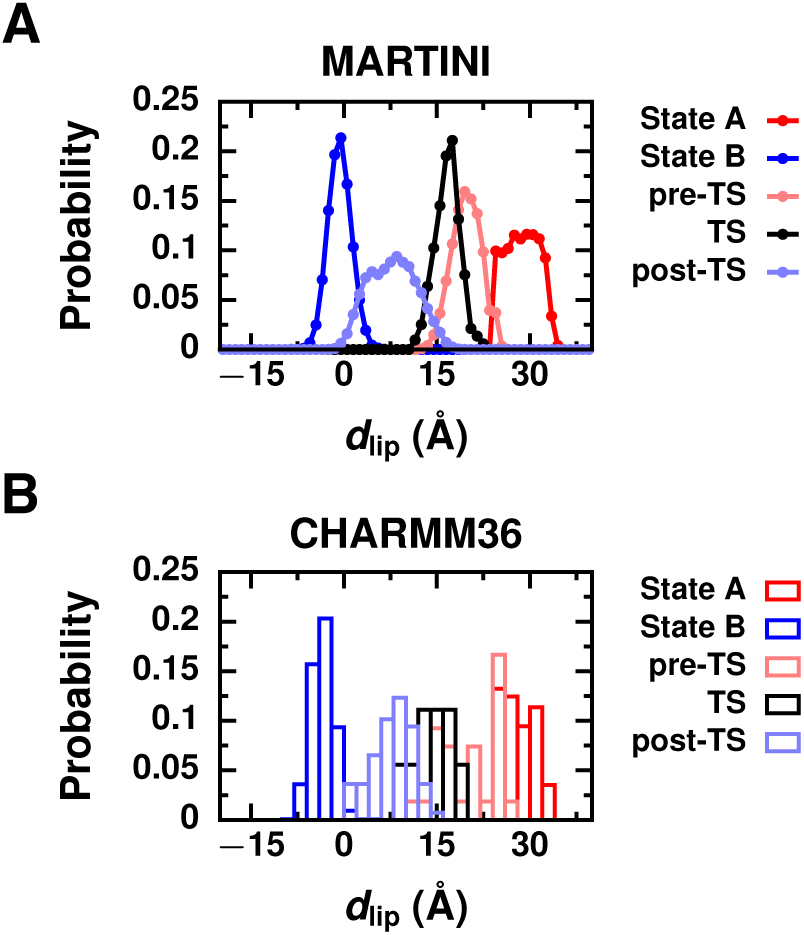
Probability distributions of *d*_lip_ from (A) MARTINI and (B) CHARMM36 simulations indicate that *d*_lip_ is 2 2 not the reaction coordinate for lipid exchange. Distributions are plotted for state A, for state B, and for three ensembles of configurations drawn from transition paths: pre-transition state (pre-TS) configurations, transition states (TS), and post-transition state (post-TS) configurations.

Based on the observation of multiple pathways for lipid insertion characterized by splayed intermediates (Figure 4), combinations of the tail displacements *d*_sn1_ and *d*_sn2_ might have been considered as pathway-specific reaction coordinates. For transitions *via* the sliding pathway, 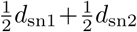 is a plausible reaction coordinate since both tails enter the bilayer at approximately the same time. For transitions *via* the sn1 splayed and sn2 splayed pathways, *d*_sn1_ and *d*_sn2_ are plausible reaction coordinates for each pathway, respectively, since the lipid is committed to fully insert after the first tail has entered the bilayer. Probability distributions of these putative reaction coordinates from MARTINI transition states are compared to the distributions from state A, state B, pre-transition state, and post-transition state configurations in Figure 8 for each pathway individually. With only 10 CHARMM36 insertion trajectories, there is insufficient data to reliably examine distributions for each pathway individually (data from all CHARMM36 transitions is shown in Figure S13). With limited overlap between the transition state distribution and distributions from all other ensembles (Figure 8),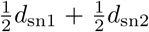, *d*_sn1_ and *d*_sn2_ appear to be potential reaction coordinates that are specific to each pathway. *d*_sn1_ cannot be used to accurately identify transition states along the sn2 splayed pathway and vice versa (Figure S14). While this indicates that transition states have values of *d*_sn1_ and *d*_sn2_ distinct from configurations in the other ensembles, it does not guarantee that 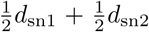, *d*_sn1_ and *d*_sn2_ are strongly correlated with *p*_B_, as required for them to be the reaction coordinates for each lipid exchange pathway.

**Figure 8:**
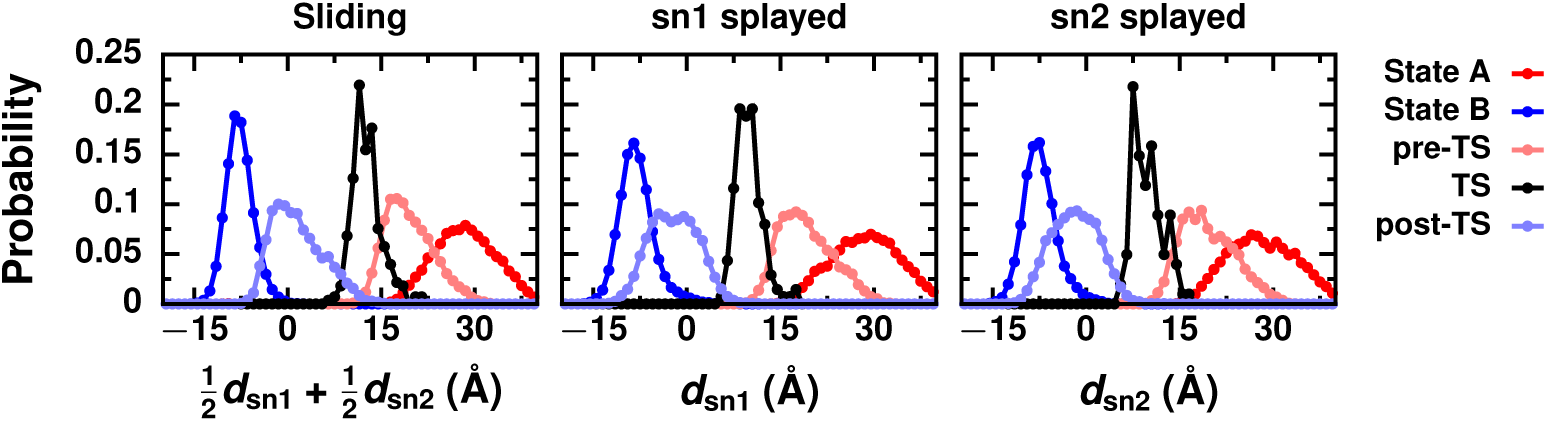
Probability distributions of 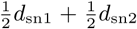, *d*_sn1_ and *d*_sn2_ from MARTINI simulations that follow a given lipid insertion pathway demonstrate that the tail displacements can reliably identify transition states. Distributions are plotte for state A, for state B, and for three ensembles of configurations drawn from transition paths: pre-transition state (pre-TS) configurations, transition states (TS), and post-transition state (post-TS) configurations.

Furthermore, if 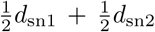 is the reaction coordinate for the sliding pathway, *d*_sn1_ for the sn1 splayed pathway, and *d*_sn2_ for the sn2 splayed pathway, then they should fully describe the dynamics of lipid exchange. In that case, the free energy surface Δ*F* (*d*_sn1_, *d*_sn2_) should have a barrier for lipid insertion and desorption to be consistent with the observed barrier crossing dynamics (Figure 4A). This free energy surface is shown in Figure 9A for the MARTINI model. Between states A (top right corner) and B (bottom left corner), slight plateaus in the free energy profile occur at combinations of *d*_sn1_ and *d*_sn2_ characteristic of splayed configurations (top left and bottom right corners). The free energy profile has a saddle point near *d*_sn1_ ≈ *d*_sn2_ ≈ 15 Å with a barrier for insertion of approximately 1 *k*_B_*T* (outlined in dashes), which is only slightly larger than the statistical error in the calculated free energy (Figure S15). This small barrier would hardly impede the dynamics of insertion. Indeed, it is significantly lower than the barrier for hydrophobic contact formation (Figure 6A), demonstrating that de creases in the tail displacments do not limit the rate of lipid insertion.

**Figure 9:**
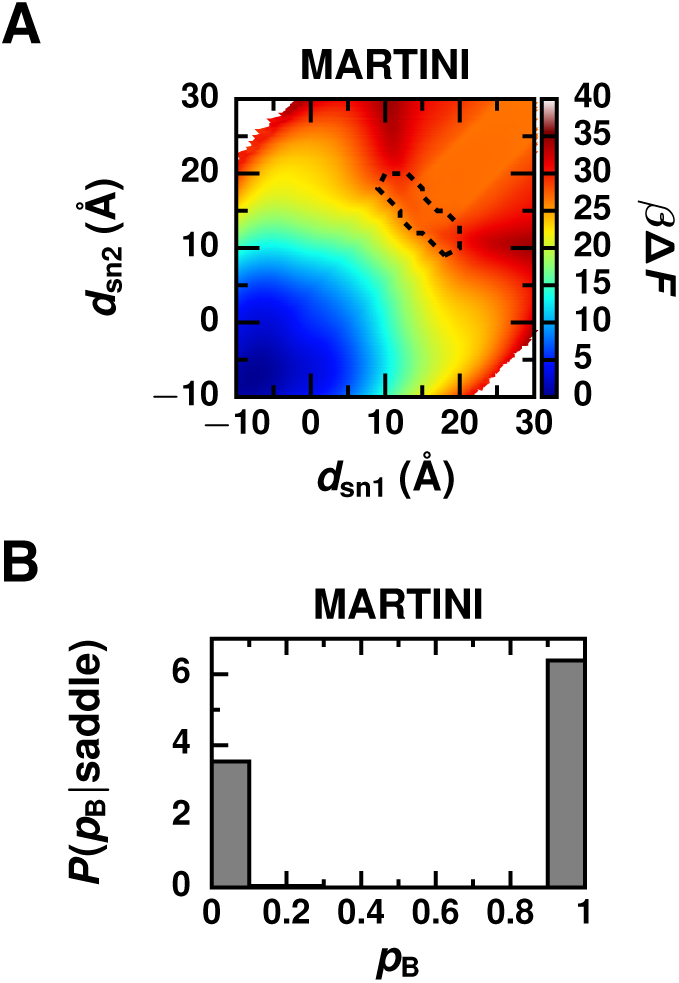
(A) Free energy surface as a function of *d*_sn1_ and *d*_sn2_ obtained from umbrella sampling simulations using the MARTINI force field lacks a significant barrier for insertion. The dashed line outlines a saddle point separating states A (top right corner) and B (bottom left corner). Statistical error in the free energy profile is shown in Figure S15. (B) Histogram of committor values, *p*_B_, for configurations located at the saddle point during umbrella sampling simulations demonstrate that the reaction coordinate is not simply a function of *d*_sn1_ and *d*_sn2_.

Finally, to determine if the tail displacements are strongly correlated with the true reaction coordinate, we constructed a histogram of *p*_B_ values for configurations found near the saddle point during umbrella sampling simulations. The histogram of *p*_B_ for configurations that are predicted to be transition states based on the free energy surface is strongly bimodal (Figure 9B). Almost all of the tested saddle point configurations are committed to state A or B, and transitions states are seldom sampled in our simulations. Thus, the lipid’s displacement normal to the bilayer, even reformulated to account for the three different insertion pathways, is not the reaction coordinate for lipid exchange.

### Calculation of Lipid Exchange Rate Enables Comparison to Experiment

Having identified the reaction coordinate, which measures hydrophobic contact formation and breakage between the lipid and membrane, we utilized this knowledge to calculate the kinetic parameters of lipid exchange. First, we calculated the lipid insertion rate, *k*_ins_, using Kramers theory (Eq. 4) as detailed in the **Methods** section and compared it to the rate obtained directly from our unbiased trajectories. Kramers theory provides an expression for the reaction rate of a process whose dynamics are diffusive and well described by an overdamped Langevin equation. ^60^ During insertion, the lipid is buffeted by solvent molecules and other membrane lipids while crossing the broad free energy barrier for hydrophobic contact formation (Figure 6A), resulting in diffusive barrier crossing dynamics (Figure S9) and making Kramers theory appropriate. The value of *k*_ins_ calculated with Kramers theory is 7.0 *µ*s^−1^ for MARTINI and 21.0 *µ*s^−1^ CHARMM36; *k*ins obtained from the mean first passage time in the spontaneous insertion simulations is 8.0 *µ*s^−1^ for MARTINI and 5.7 *µ*s^−1^ for CHARMM36. The insertion rates calculated based on hydrophobic contact formation differ from those obtained from dynamical simulations by a factor of 4 or less. Such good agreement demonstrates that hydrophobic contacts describe the dynamics of lipid insertion, an elementary step of lipid exchange, with quantitative accuracy. Unfortunately, the timescale of insertion is too fast to measure with straightforward experimental methods.

To compare our results to experiments, we sought to calculate instead the much slower lipid exchange rate, *k*_ex_. Although we obtained *k*_ins_ directly from simulations, *k*_ex_ cannot be obtained in the same manner. Complete exchange events, which involve lipid desorption and diffusion to another membrane in addition to insertion, could not be simulated due to their prohibitively long waiting times. Given the good agreement between *k*_ins_ calculated directly from simulation and from Kramers theory, we instead calculated *k*_ex_ from a Smoluchowski equation that contains information about the free energetics of hydrophobic lipid–membrane contacts.

To experimentally probe passive lipid exchange, two populations of vesicles, one initially composed of labeled lipids and the other initially of unlabeled ones, are combined in solution. *k*_ex_ is then determined by monitoring the rate of mixing of labeled lipids between these two populations of vesicles. ^18,26,27,75–77^ We can obtain an expression for the rate of mixing by first considering how the concentration of labeled lipids in a single vesicle changes over time. The concentration of labeled lipids in each vesicle will adopt a steady state with the concentration in the solution surrounding that vesicle. At steady state, the flux of lipids over any closed surface around a vesicle is constant. Importantly, since lipids that desorb may rapidly return to their original vesicle given the small barrier for insertion compared to desorption (Figure 6A), the flux contains contributions from lipids both leaving and entering a vesicle. According to the Smoluchowski equation, the flux, *J*, of lipids radially into a vesicle is

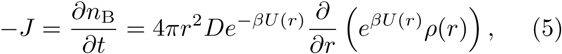

where *n*_B_ is the number of labeled lipids in a vesicle, *r* is the radial distance from the center of a vesicle, *D* is the diffusion coefficient of a lipid in solution, *U* (*r*) is the effective interaction potential between a lipid and a vesicle, and *ρ*(*r*) is the concentration of labeled lipids at a distance *r*. As for the calculations using Kramers theory, we use min(*d*_CC_) to describe interactions between a lipid and a vesicle in 3D space. *U* (*r*) is then the free energy profile as a function of min(*d*_CC_) (Figure S10). We set *U* (*r*) = 0 far away from a vesicle and shift the peak of the free energy barrier to *r* = 50 nm, a typical radius of large unilamellar vesicles (LUVs) used in experimental studies. ^18,26,27,75–77^ Assuming that the concentration profile of labeled lipids reaches steady state much faster than *n*_*B*_ is varying, the time dependence of the bulk concentration of labeled lipids, 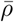, can be regarded as constant while calculating the steady-state profile 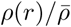. Assuming as well that equilibration within state B is fast, the concentration of labeled lipids within a vesicle obeys equilibrium statistics,

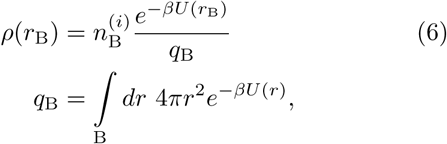

where *i* indicates whether the vesicle was initially composed of labeled (*i* = 1) or unlabeled (*i* = 2) lipids, *r*_B_ is a particular location within a vesicle, *q*_B_ is the partition function for labeled lipids in a vesicle, and the integral is performed over the range of *r* designated as state B. With these two assumptions, Eq. 5 integrates to

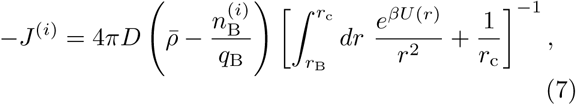

where *r*_c_ is the distance beyond which *U* (*r*) is zero. Finally, the rate of mixing is obtained by considering how the difference between the number of labeled lipids in type 1 vesicles and in type 2 changes over time:

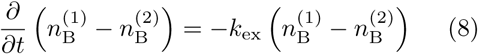

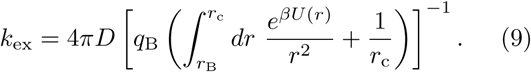

Eq. 8 correctly reflects the fact that lipid exchange is a first order process. ^17–19,21,24–31^ Both the first term in the denominator of Eq. 9, which dominates when the free energy barrier is very large, and the second term, which dominates when diffusion is rate limiting, are key to the rate of lipid exchange.

The values of *k*_ex_ calculated with Eq. 9 for MARTINI DLPC and CHARMM36 DMPC are compared to experimental values in Table 1. In the case of MARTINI DLPC, our calculated *k*_ex_ is in agreement with experimental values. However, in the case of CHARMM36 DMPC, our calculated *k*_ex_ differs from experimental values by six orders of magnitude. Some discrepancy between experiment and simulation/theory is to be expected because small errors in the free energy profile are magnified exponentially in *k*_ex_. Yet, a six-order-of-magnitude difference between theoretical and experimental rates corresponds to a significant underestimation of the free energy barrier of approximately 12 *k*_B_*T* for CHARMM36. This large discrepancy could be due to any number of the following reasons:

**Table 1:**
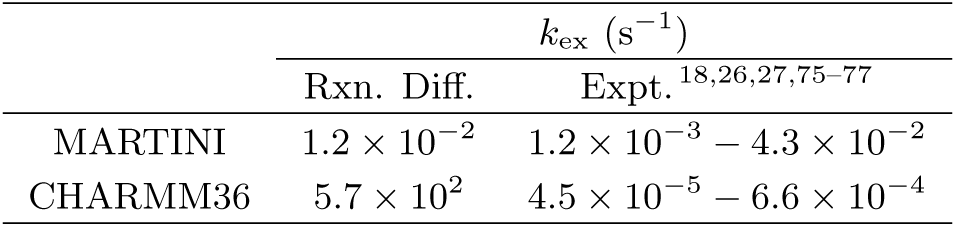
Rates of lipid exchange.

1. Sampling errors, which are within about 1 *k*_B_*T* throughout the free energy surface (Figure S9), may influence our rate calculations.
2. Inaccuracies in the force field may cause errors in the calculated free energy surface. Based on reported differences between permeabilities and partition coefficients for water, alkanes, and other small molecules obtained from simulations with CHARMM36 and from experiments, ^78,79^ we estimate that errors due to the force field are approximately 1 −2 *k*_B_*T*. We also repeated our calculations using a different force field (Stockholm lipids, also known as Slipids ^80,81^) as detailed in the SI. The height of the free energy barrier is approximately 2 *k*_B_*T* higher for Slipids compared to CHARMM36 (Figure S16), resulting in an estimate of *k*_ex_ that still differs from experiment by about five orders of magnitude.
3. The ionic strength of the solution may influence the free energy surface. Our simulations are performed with neat water, but experimental systems are conducted in buffer, which contains significant amounts of monovalent salts, such as NaCl or KCl. ^18,75–77^ The addition of NaCl or KCl could salt out the free lipids in solution, increasing the difference in free energy between state A and B. Additionally, monovalent cations bind to the carbonyl region of phosphatidylcholine membranes, causing the thickness of the bilayer and order of the tails to increase. ^82–84^ These structural changes are expected to increase the free energy barrier for hydrophobic contact formation.
4. The assumptions we made to obtain Eq. 7 may not be consistent with what occurs in experimental systems, causing discrepancies between our calculated *k*_ex_ and experimental measurements. In writing down a Smoluchowski equation (Eq. 5), we assumed that lipids exchange between stable, spherical vesicles of uniform size. This may be justified since vesicles composed of biological phospholipids are kinetically-trapped, metastable states, which by definition do not quickly relax into equilibrium lamellar structures. ^85–88^ But, we lack the requisite knowledge about the processes and associated timescales by which vesicles relax into equilibrium structures to confirm the validity of this assumption. Other relaxation processes, such as uncatalyzed vesicle fusion, which has an experimentally measured rate similar to lipid exchange, ^89^ could occur simultaneously. We note that lipid flip-flop, the process by which a lipid moves between leaflets of the same bilayer, ^1,5^ also has a rate measured under some experimental conditions that is similar to the exchange rate.^75,90^ Unlike vesicle fusion, lipid flip-flop is accounted for in experimental determination of lipid exchange rates.
5. Similar issues of random error and unjustified kinetic assumptions could plague the inference of exchange rates from laboratory measurements.

## CONCLUSIONS

The breakage of hydrophobic contacts limits the rate of passive lipid transport. To reach this conclusion, we investigated the elementary steps of lipid exchange, lipid insertion into and desorption from a membrane, using molecular dynamics simulations of the coarse-grained MARTINI and all-atom CHARMM36 lipid models. Results from MARTINI and CHARMM36 provide consistent pictures; even with a coarse description of lipids and water, the MARTINI model captures the essential features of lipid exchange exhibited in an allatom model. We discovered that the reaction coordinate for passive lipid exchange measures the formation and breakage of hydrophobic lipid–membrane contacts, which gives rise to a free energy barrier for both lipid desorption and insertion.

Thus, knowledge of the reaction coordinate resolves previous qualitative discrepancies between simulations, which predicted that there is no barrier for lipid insertion, and experiments, which indicated that there is a barrier. This barrier likely plays an important biological role: A barrier for lipid insertion ensures that membrane compositions, which are spateotemporally regulated to maintain cell homeostasis, ^1,2^ are not easily disrupted. We suspect that the formation of hydrophobic contacts may generally give rise to a free energy barrier for transporting amphiphiles, including synthetic surfactants ^91^ and lipopeptides, ^55,92^ which is not resolved by monitoring the amphiphile’s center-of-mass displacement from a membrane.

Additionally, knowledge of the reaction coordinate allowed us to formulate a Smoluchowski equation to model lipid exchange between vesicles, which occurs over time and length scales inaccessible in MD simulations, and calculate the rate of lipid exchange. Differences between our calculated lipid exchange rate and experimental measurements indicate that considerable quantitative discrepancies between simulation and experiment still exist. Future studies will be performed to better assess the sources of these discrepancies.

Finally, this knowledge provides a foundation to understand how catalysts of lipid exchange work at a molecular level. Lipid transfer proteins may efficiently extract lipids from membranes by lowering the activation free energy barrier for hydrophobic contact breakage. Interestingly, catalysts of vesicle fusion, the key step in vesicular lipid transport, may function in a similar way. For example, carbon nanotubes aid vesicle fusion by facilitating the formation of hydrophobic contacts between two vesicles, ^93^ and viral fusion peptides are thought to catalyze fusion by promoting hydrophobic lipid tail protrusions and contacts.^61^ Thus, common physical properties of lipids, specifically their hydrophobicity, may be exploited *in vivo* to precisely control both non-vesicular and vesicular lipid transport.

## Acknowledgement

J.R.R. acknowledges the support of the National Science Foundation Graduate Research Fellowship Program under Grant No. DGE 1752814. P.L.G. was supported by the Director, Office of Basic Energy Sciences, Office of Science, US Department of Energy under Contract DE-AC02-05CH11231, through the Chemical Sciences Division of Lawrence Berkeley National Laboratory. This work used the Extreme Science and Engineering Discovery Environment (XSEDE), which is supported by National Science Foundation grant number ACI-1548562; specifically, it utilized XSEDE Comet at the San Diego Supercomputing Center through allocation CHE180038. We thank Prof. Baron Peters for providing us with maximum likelihood approach code. We thank Dr. Georg Menzl and Layne Frechette for valuable discussions and helpful comments.

## Supporting Information Available

Descriptions of all order parameters used to analyze transition paths; descriptions of simulations with the Slipids force field; description and results of maximum likelihood approach to identify a reaction coordinate; demonstration of equilibration of initial lipid bilayers; probability distributions used to determine stable state definitions; probability distributions of additional order parameters; statistical errors in computed free energy surfaces; minimum free energy paths along min(*d*_CC_) and *n*_CC_; 1D free energy profiles of min(*d*_CC_) and *n*_CC_; results from simulations with the Slipids force field; harmonic bias parameters used for umbrella sampling simulations; frequency of each insertion pathway; movies of lipid insertion trajectories.

## Graphical TOC Entry

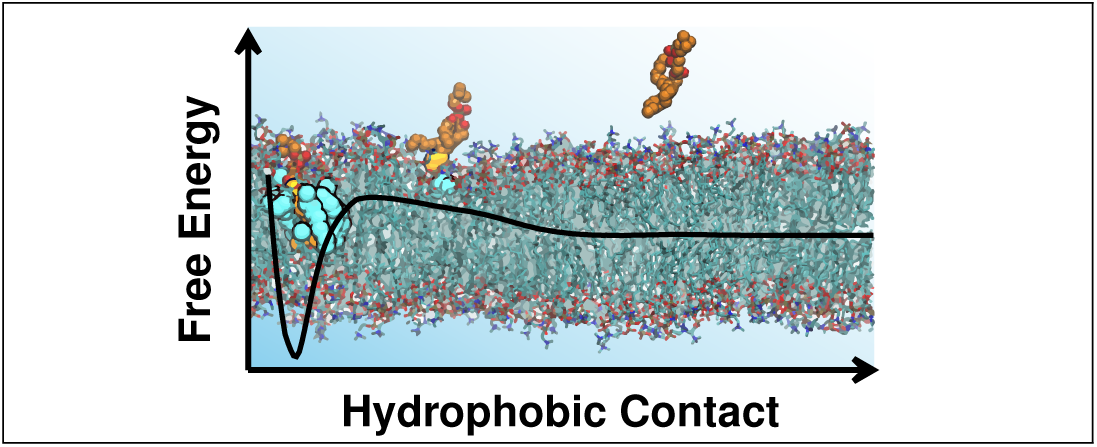

