## Supplemental Information for "Breakage of Hydrophobic Contacts Limits the Rate of Passive Lipid Exchange Between Membranes"

### SUPPLEMENTARY METHODS

**Order Parameters Used to Characterize Transition Paths.** In addition to the order parameters described in the main text, $d_{\text{lip}}$, $d_{\text{sn1}}$, $d_{\text{sn2}}$, $\min(d_{\text{CC}})$, and $n_{\text{CC}}$, we evaluated additional order parameters for their potential as reaction coordinates. They are as follows:

$n_{\text{w,X}}^{(\text{lip})}$ The number of water molecules within the first solvation shell of bead (atom) X of the tagged MARTINI (CHARMM36) lipid. Water beads (water molecules with oxygen atoms) within a distance $d_{\text{solv}}^{(\text{lip})}(\text{X})$ of X were counted. $d_{\text{solv}}^{(\text{lip})}(\text{X})$ is the location of the first minimum of the radial distribution function between water beads (oxygens) and X calculated from a simulation of a single lipid in solution.

$n_{\text{w,X}}^{(\text{mem})}$ The number of water molecules within the first solvation shell of bead (atom) X of MARTINI (CHARMM36) lipids in the membrane that are nearby the tagged lipid. Only lipids in the membrane whose distance to the tagged lipid in the plane parallel to the bilayer surface ($xy$ plane) was within 8 Å were considered. Water beads (water molecules with oxygen atoms) within a distance $d_{\text{solv}}^{(\text{mem})}(\text{X})$ of X were counted. $d_{\text{solv}}^{(\text{mem})}(\text{X})$ is the location of the first minimum of the radial distribution function between water beads (oxygens) and X calculated from a simulation of an unperturbed lipid bilayer. The value of $d_{\text{solv}}^{(\text{mem})}(\text{X})$ for beads C1A and C1B (hydrophobic carbons closest to the glycerol group) were used for all hydrophobic carbons in the lipid tails.

$n_{\text{X,Y}}^{(\text{cyl})}$ The number of X within a cylinder extending in $z$ from Y to the midplane of the bilayer. X is either water beads (oxygens) with label X = w, all lipid beads (atoms) with label X = lip, hydrophilic lipid beads (atoms) with label X = philic, hydrophobic lipid beads (atoms) with label X = phobic, or all beads (atoms) with label X = all. Y is either the center-of-mass (COM) of the tagged lipid with label Y = lip, the terminal carbon of the sn1 tail of the tagged lipid with label Y = sn1, or the terminal carbon of the sn2 tail of the tagged lipid with label Y = sn2. For Y = lip, the radius of the cylinder is 8 Å; for Y = sn1, the radius is $d_{\text{solv}}^{(\text{lip})}(\text{C3B})$ for MARTINI and $d_{\text{solv}}^{(\text{lip})}(\text{C314})$ for CHARMM36; for Y = sn2 the radius is $d_{\text{solv}}^{(\text{lip})}(\text{C3A})$ for MARTINI and $d_{\text{solv}}^{(\text{lip})}(\text{C214})$ for CHARMM36.

$\rho_{\text{X,Y}}^{(\text{cyl})}$ The number density of X within a cylinder extending in $z$ from Y to the midplane of the bilayer, where X = w, lip, philic, phobic, or all and Y = lip, sn1, or sn2. The radius of the cylinder depends on Y just as for $n_{\text{X,Y}}^{(\text{cyl})}$.

$\langle \delta z_{\text{phos}} \rangle_{\text{mem}}$ Average height fluctuation of lipids in the membrane that are within 8 Å of the tagged lipid in the $xy$ plane. $\delta z_{\text{phos}} = z_{\text{phos}} - \langle z_{\text{phos}} \rangle$ is the deviation of the $z$ position of the phosphate group of a lipid nearby the tagged lipid from the average $z$ position of the phosphate group of all lipids in the leaflet.

$\langle n_{\text{ET}} \rangle_{\text{mem}}$ Average number of exposed lipid tails flipped out of the bilayer. An exposed tail has at least one hydrophobic carbon bead (atom) with a $z$ position greater than $\langle z_{\text{phos}} \rangle$. Only lipids nearby the tagged lipid, which are within 8 Å of the tagged lipid in the $xy$ plane, were considered.

$\min(d_{\text{HT,Y}})$ The minimum distance between head groups of lipids in the membrane and the terminal carbon of one tail of the tagged lipid in the $xy$ plane. Specifically, $d_{\text{HT,Y}}$ is measured from bead NC3 (atom N) to either terminal carbon of the tagged lipid's tails, labeled Y = sn1 or sn2, for MARTINI (CHARMM36).

$A_{\mathrm{defect,Y}}$ Area of packing defects in the membrane near one tail of the tagged lipid, labeled Y = sn1 or sn2. Defects that expose hydrophobic portions of the membrane were identified with PackMem[1,2] on a 1 Å × 1 Å grid. The tagged lipid was excluded in the analysis to identify defects. A defect is considered near Y if any grid point that makes up the defect is within 1 Å of Y in the $xy$ plane.

$E_{\mathrm{int}}$ Interaction energy between the tagged lipid and the rest of the system.

$\min(d_{\mathrm{CC,Y}})$ Hydrophobic contacts between one tail of the tagged lipid and membrane. $d_{\mathrm{CC,Y}}$ is the distance between a hydrophobic carbon in the sn1 tail, labeled Y = sn1, or sn2 tail, labeled Y = sn2, of the tagged lipid and of the closest leaflet.

$n_{\mathrm{CC,Y}}$ Number of close hydrophobic contacts between one tail of the tagged lipid, labeled Y = sn1 or sn2, and closest leaflet. Any pair of hydrophobic carbons with a distance $d_{\mathrm{CC}} \leq 14$ Å for MARTINI lipids or with $d_{\mathrm{CC}} \leq 10$ Å for CHARMM36 lipids were counted as close contacts.

**Stockholm Lipids (Slipids) Simulations.** *Molecular Dynamics Simulations.* To simulate a bilayer of 128 Slipids[3,4] DMPC lipids for use in subsequent umbrella sampling (US) simulations, we used a protocol similar to that used for CHARMM36 lipids. First, the bilayer was energy minimized prior to undergoing a two-stage equilibration at 320 K and 1 bar. The first 250 ps equilibration utilized the Berendsen barostat[5] for semi-isotropic pressure coupling with a coupling time constant of 2 ps and isothermal compressibility of $4.5 \times 10^{-5}$ bar$^{-1}$, and the second 250 ps equilibration utilized the Parinello-Rahman barostat[6] with a coupling time constant of 10 ps. Then, a 50 ns production run was performed to allow the bilayer to fully equilibrate (Figure S1). The lipids and solvent were coupled to separate Nosé-Hoover thermostats[7,8] using a coupling time constant of 1 ps to maintain the temperature. Dynamics were evolved according to the leapfrog algorithm[9] using a 2 fs time step. All bonds were constrained using the LINCS algorithm.[10] Lennard-Jones forces were smoothly switched off between 1.4 and 1.5 nm. Coulomb interactions were truncated at 1.5 nm, and long-ranged Coulomb interactions were calculated using Particle Mesh Ewald (PME) summation[11] with a Fourier spacing of 0.12 nm and an interpolation order of 4. A long-range analytic dispersion correction was applied to both energy and pressure to account for the truncation of Lennard-Jones interactions. Neighbor lists were constructed with the Verlet algorithm.[12]

Next, a system with a tagged lipid in solution was built for use in US simulations. This system was energy minimized and equilibrated using the protocol described for the Slipids bilayer system with the addition of harmonic restraints on the $z$ coordinates of all heavy atoms of the tagged lipid.

*Free Energy Calculations.* To obtain the 2D free energy surface $\Delta F(\min(d_{\mathrm{CC}}), n_{\mathrm{CC}})$ for the Slipids system using US simulations, the same number of windows and the same harmonic biases on $\min(d_{\mathrm{CC}})$ and $n_{\mathrm{CC}}$ were used as for the CHARMM36 system (Table S2). To generate initial configurations for each window, a 20 ns steered molecular dynamics (SMD) simulation was performed. During the SMD simulation, the system was pulled along the minimum free energy path determined for CHARMM36 (Figure S9) using a harmonic bias on $\min(d_{\mathrm{CC}})$ with a spring constant of 5,000 kJ/mol/nm$^2$ and on $n_{\mathrm{CC}}$ with a spring constant of 0.005 kJ/mol/contact$^2$. Each US window was initialized with a configuration obtained from the SMD simulation that has a value of ($\min(d_{\mathrm{CC}})$, $n_{\mathrm{CC}}$) close to the center of the window's bias. Each window was simulated for 24 ns. The first 4 ns of data from these windows was discarded to account for equilibration. Finally, data from all windows was combined with the weighted histogram analysis method (WHAM)[13] to obtain a free energy surface as a function of $\min(d_{\mathrm{CC}})$ and $n_{\mathrm{CC}}$. Error bars were calculated as the standard error of free energy surfaces estimated from five independent 5 ns blocks.

**Maximum Likelihood Approach to Identify a Reaction Coordinate.** The maximum likelihood approach[14,15] identifies a reaction coordinate $r_c$, which is a linear combination of order parameters $q_i$, that best represents the outcomes of individual committor calculations when inserted into a model for the committor. The model for the committor is

$$p_{\mathrm{B}}(r_c(\boldsymbol{q})) = \frac{1 + \tanh(r_c(\boldsymbol{q}))}{2}.$$

Specifically, the form of the reaction coordinate is

$$r_c(\boldsymbol{q}) = \alpha_0 + \sum_{i=1}^{M} \alpha_i q_i,$$

where $M$ is the number of order parameters used to construct $r_c$. All order parameters $q_i$ are scaled to the range [0,1], and their units are absorbed into the coefficients $\alpha_i$. The coefficient $\alpha_0$ is included as an offset so that $r_c = 0$ for transition states. The coefficients $\alpha_i$ are determined by maximizing the likelihood

$$\mathcal{L} = \prod_{k=1}^{N_{\mathrm{B}}} p_{\mathrm{B}}(r_c(\boldsymbol{q})) \prod_{k=1}^{N_{\mathrm{A}}} \left[1 - p_{\mathrm{B}}(r_c(\boldsymbol{q}))\right],$$

where $N_{\mathrm{B}}$ is the number of trials used in the committor calculations that end in state B and $N_{\mathrm{A}}$ is the number of trials that end in state A. As $M$ is increased, the number of fitting parameters $\alpha_i$ increases, resulting in an improved reaction coordinate and increased likelihood at the cost of physical insight. A Bayesian information criterion can be used to determine when including an additional order parameter results in a significantly improved reaction coordinate. For $N = N_{\mathrm{A}} + N_{\mathrm{B}}$ realizations of the likelihood, a reaction coordinate composed of $M+1$ order parameters provides a significant improvement over a reaction coordinate composed of $M$ order parameters if $\ln \mathcal{L}$ increases by more than the Bayesian information criterion, $\frac{1}{2} \ln N$.

We used the maximum likelihood approach to construct $r_c^{(M=48)}$ as a linear combination of up to 48 order parameters, excluding $\min(d_{\mathrm{CC}})$ and $n_{\mathrm{CC}}$, for the MARTINI model. Additionally, we used the maximum likelihood approach to construct $r_c$ from order parameters including $\min(d_{\mathrm{CC}})$ and $n_{\mathrm{CC}}$ for both MARTINI and CHARMM36 models. $r_c$ constructed as a linear combination of $\min(d_{\mathrm{CC}})$ and $n_{\mathrm{CC}}$ was used to identify dividing surfaces between states A and B where $r_c = 0$. For MARTINI, the likelihood for each tested reaction coordinate was evaluated on $N = 4,569,150$ commitor trials found along transition paths, corresponding to a Bayesian information criterion of $\frac{1}{2} \ln N = 7.667$. For CHARMM36, the likelihood for each tested reaction coordinate was evaluated based on $N = 2,760$ commitor trials found along transition paths, corresponding to a Bayesian information criterion of $\frac{1}{2} \ln N = 3.961$.

### SUPPLEMENTARY FIGURES

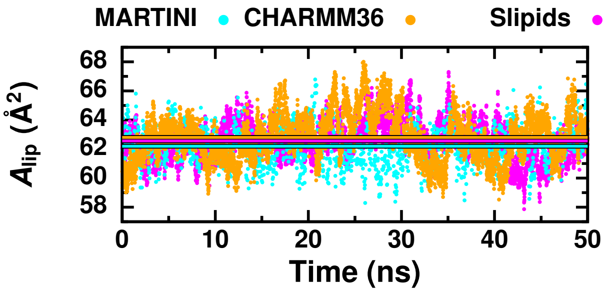


**Figure S1. Equilibration of initial lipid bilayer.** Area per lipid, $A_{\mathrm{lip}}$, during MD simulations of a lipid bilayer composed of 128 lipids. The straight lines that are outlined in black indicate the average area per lipid during the MARTINI simulation (cyan), CHARMM36 simulation (orage), and Slipids simulation (magenta).

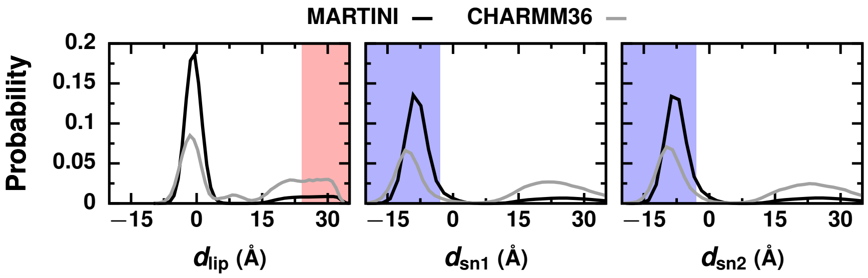


**Figure S2. Determination of stable state definitions.** Probability distributions of $d_{\text{lip}}$, $d_{\text{sn1}}$, and $d_{\text{sn2}}$ from MARTINI and CHARMM36 spontaneous insertion trajectories. State A configurations, in which the tagged lipid is fully solvated, are located in the red region. State B configurations, in which the tagged lipid is in the bilayer, are located in the blue region. The small peak around $d_{\text{lip}} = 20$ Å arises from configurations in which the tagged lipid is absorbed to the bilayer. The peak in the distribution from CHARMM36 simulations around $d_{\text{lip}} = 8$ Å arises from splayed lipid configurations.

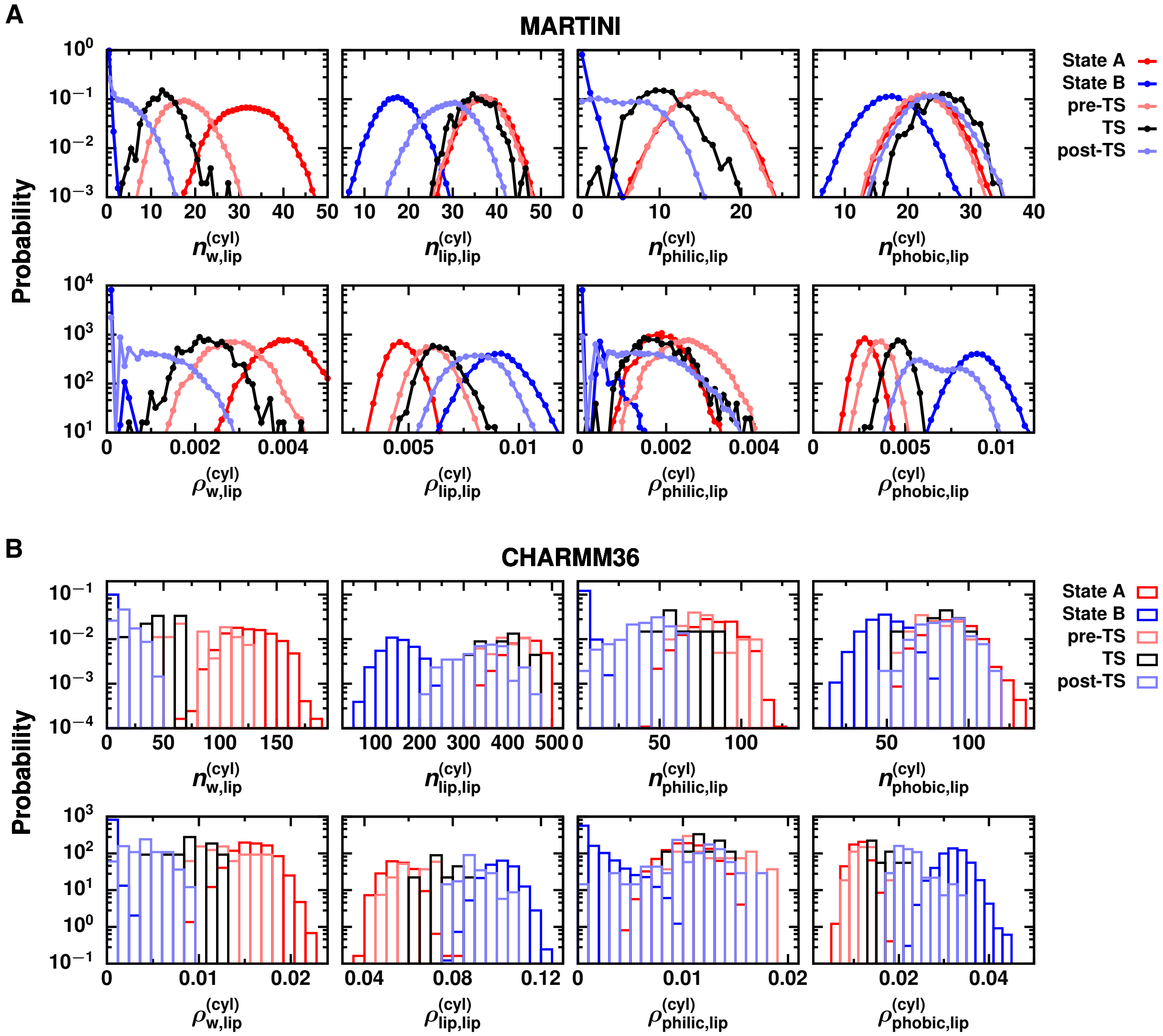


**Figure S3. Cavities around the tagged lipid are not observed during insertion. The number of water molecules and hydrophilic lipid fragments below the tagged lipid decreases during insertion while the density of hydrophobic lipid fragments increases.** Probability distributions of the number, $n^{(\mathrm{cyl})}_{\mathrm{X,lip}}$, and number density, $\rho^{(\mathrm{cyl})}_{\mathrm{X,lip}}$, of water and different types of molecular fragments between the tagged lipid and midplane of the bilayer for (A) MARTINI and (B) CHARMM36 simulations. Distributions are plotted for state A, for state B, and for three ensembles of configurations drawn from transition paths: pre-transition state (pre-TS) configurations identified by $p_{\mathrm{B}} = 0$, transition states (TS) identified by $0.45 \leq p_{\mathrm{B}} \leq 0.55$ for MARTINI and by $0.4 \leq p_{\mathrm{B}} \leq 0.6$ for CHARMM36, and post-transition state (post-TS) configurations identified by $p_{\mathrm{B}} = 1$.

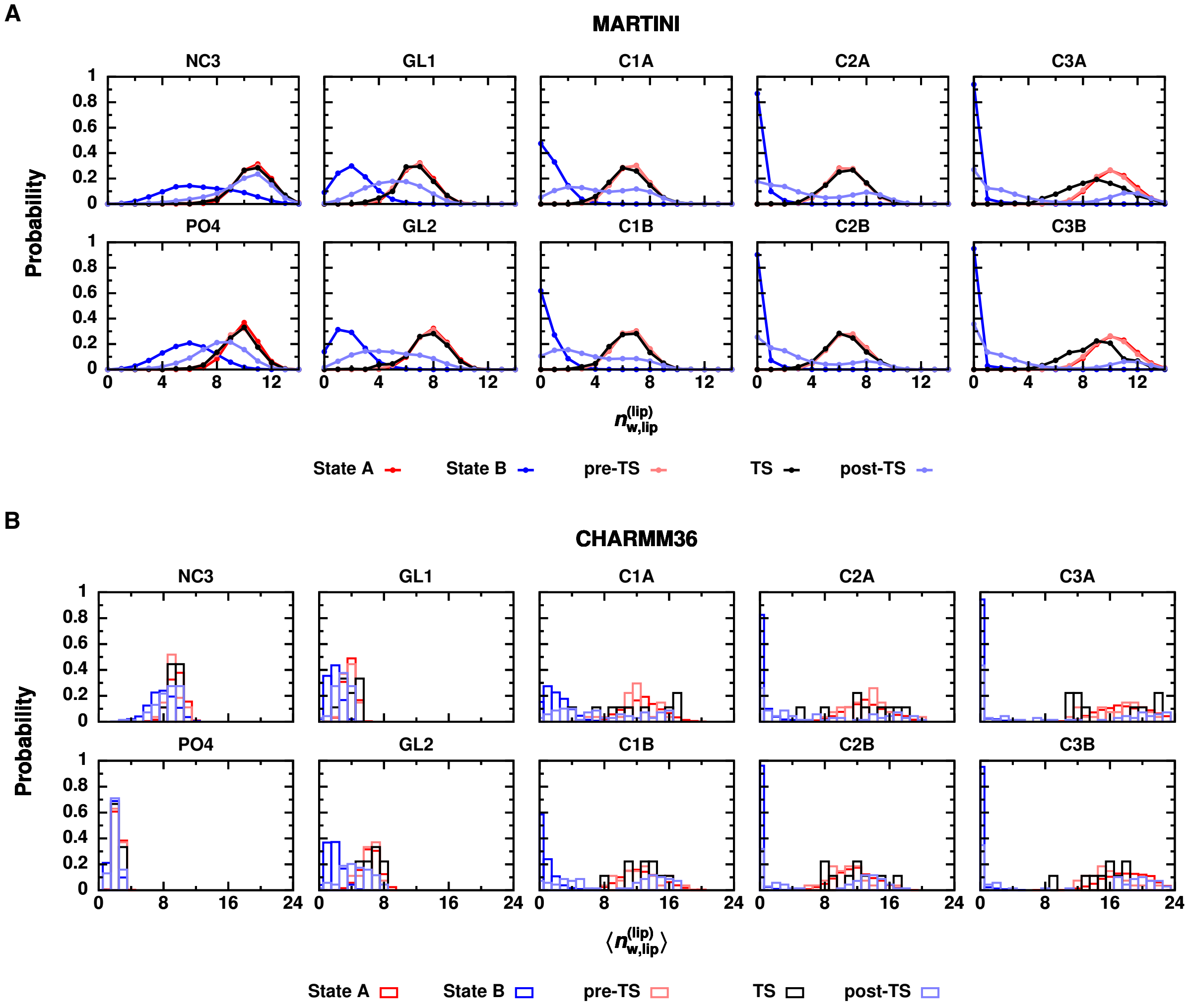


**Figure S4. Water molecules are shed from the tagged lipid's first solvation shell during insertion.** Probability distributions of the number of water molecules within the first solvation shell of the tagged lipid, $n_{\mathrm{w,X}}^{(\mathrm{lip})}$. (A) For MARTINI, the number of water beads around each bead of the tagged lipid is shown. (B) For CHARMM36, $n_{\mathrm{w,X}}^{(\mathrm{lip})}$ is averaged over the atoms that map to a given MARTINI bead: Atoms N, C13, C14, C15, C12, and C11 are mapped to bead NC3; atoms P, O13, O14, O11, and O12 are mapped to bead PO4; atoms C1, C2, O21, C21, O22, and C22 are mapped to bead GL1; atoms C3, O31, C31, O32, and C32 are mapped to bead GL2; atoms C23, C24, C25, and C26 are mapped to bead C1A; atoms C27, C28, C29, and C210 are mapped to bead C2A; atoms C211, C212, C213, and C214 are mapped to bead C3A; atoms C33, C34, C35, and C36 are mapped to bead C1B; atoms C37, C38, C39, and C310 are mapped to bead C2B; atoms C311, C312, C313, and C314 are mapped to bead C3B. Beads C1B, C2B, and C3B comprise the sn1 tail, and beads C1A, C2A, and C3A comprise the sn2 tail. Distributions are plotted for state A, for state B, and for three ensembles of configurations drawn from transition paths: pre-transition state (pre-TS) configurations identified by $p_{\mathrm{B}} = 0$, transition states (TS) identified by $0.45 \leq p_{\mathrm{B}} \leq 0.55$ for MARTINI and by $0.4 \leq p_{\mathrm{B}} \leq 0.6$ for CHARMM36, and post-transition state (post-TS) configurations identified by $p_{\mathrm{B}} = 1$.

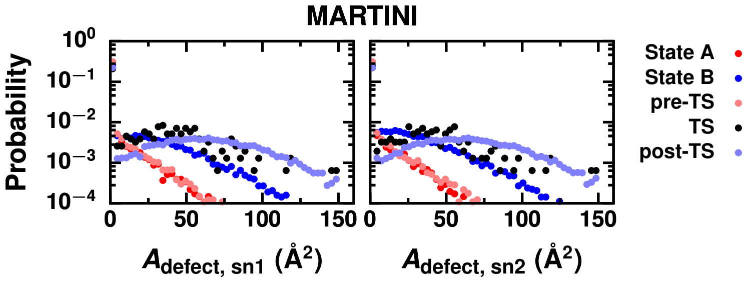


**Figure S5. Membrane packing defects that expose hydrophobic lipid fragments are increased at the transition state and when the lipid is committed to entering the membrane.** Probability distributions of the area of defects located nearby either tail of the tagged lipid in the $xy$ plane, $A_{\text{defect,Y}}$. Distributions are plotted for state A, for state B, and for three ensembles of configurations drawn from transition paths: pre-transition state (pre-TS) configurations identified by $p_B = 0$, transition states (TS) identified by $0.45 \le p_B \le 0.55$ for MARTINI and by $0.4 \le p_B \le 0.6$ for CHARMM36, and post-transition state (post-TS) configurations identified by $p_B = 1$. Becuase the tagged lipid is excluded from the analysis of defects, larger defects are observed in state B than in state A; these defects correspond to the tagged lipid.

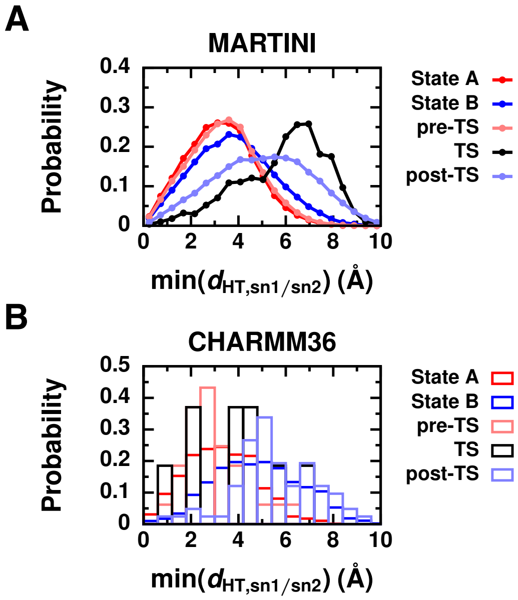


**Figure S6. The distance between the tagged lipid's tails and hydrophilic head groups of membrane lipids is increased at the transition state.** Probability distributions of the minimum distance between the head groups of the membrane lipids and either tail of the tagged lipid measured in the $xy$ plane, $\min(d_{\text{HT,sn1/sn2}})$, from (A) MARTINI and (B) CHARMM36 simulations. Distributions are plotted for state A, for state B, and for three ensembles of configurations drawn from transition paths: pre-transition state (pre-TS) configurations identified by $p_B = 0$, transition states (TS) identified by $0.45 \le p_B \le 0.55$ for MARTINI and by $0.4 \le p_B \le 0.6$ for CHARMM36, and post-transition state (post-TS) configurations identified by $p_B = 1$.

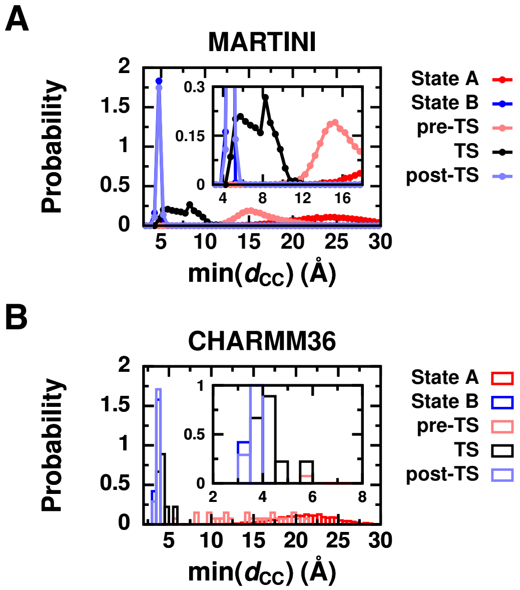


**Figure S7. The minimum distance between hydrophobic lipid and membrane carbons may be the reaction coordinate.** Probability distributions of the min($d_{\mathrm{CC}}$) from (A) MARTINI and (B) CHARMM36 simulations. Distributions are plotted for state A, for state B, and for three ensembles of configurations drawn from transition paths: pre-transition state (pre-TS) configurations identified by $p_{\mathrm{B}} = 0$, transition states (TS) identified by $0.45 \leq p_{\mathrm{B}} \leq 0.55$ for MARTINI and by $0.4 \leq p_{\mathrm{B}} \leq 0.6$ for CHARMM36, and post-transition state (post-TS) configurations identified by $p_{\mathrm{B}} = 1$. Magnified views of the transition state distributions are shown in the insets.

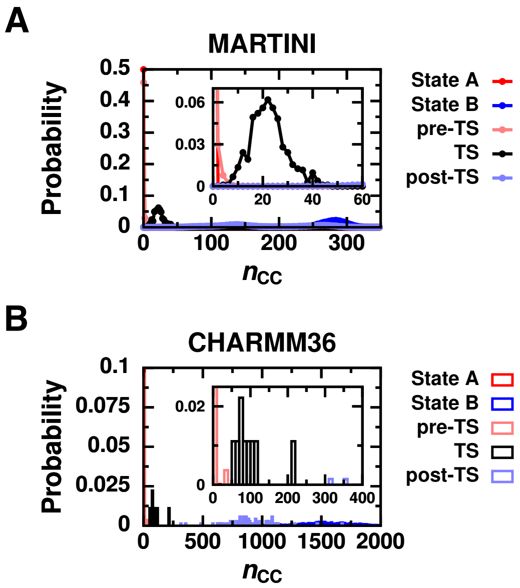


**Figure S8. The number of close hydrophobic lipid–membrane contacts may be the reaction coordinate.** Probability distributions of $n_{\mathrm{CC}}$ from (A) MARTINI and (B) CHARMM36 simulations. Distributions are plotted for state A, for state B, and for three ensembles of configurations drawn from transition paths: pre-transition state (pre-TS) configurations identified by $p_{\mathrm{B}} = 0$, transition states (TS) identified by $0.45 \leq p_{\mathrm{B}} \leq 0.55$ for MARTINI and by $0.4 \leq p_{\mathrm{B}} \leq 0.6$ for CHARMM36, and post-transition state (post-TS) configurations identified by $p_{\mathrm{B}} = 1$. Magnified views of the transition state distributions are shown in the insets.

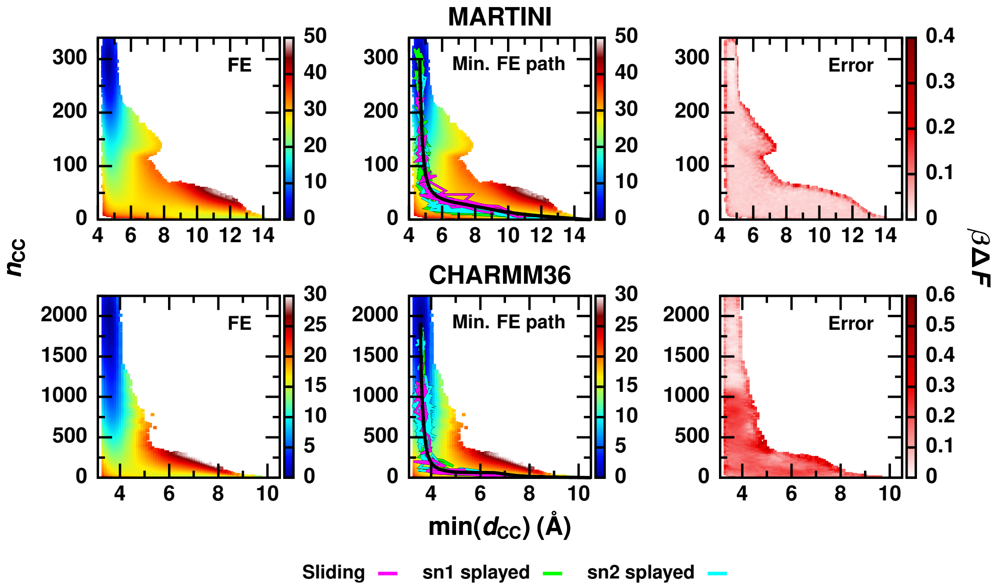


**Figure S9. Transition paths follow the minimum free energy path in the space of min($d_{CC}$) and $n_{CC}$.** (Left panel) Free energy surfaces as a function of min($d_{CC}$) and $n_{CC}$ obtained from umbrella sampling simulations. (Middle panel) Example transitions paths for each lipid insertion pathway are plotted along with the minimum free energy path (black line). (Right panel) Block standard error of the free energy surface calculated from five simulation blocks.

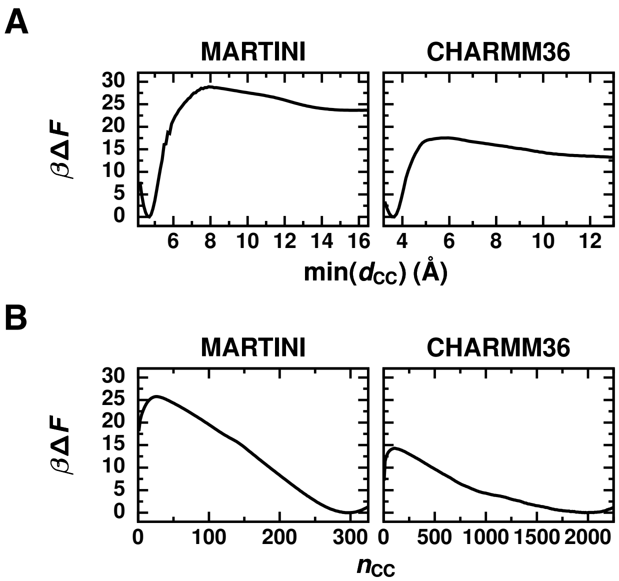


**Figure S10. A free energy barrier for lipid insertion exists due to hydrophobic lipid–membrane contact formation.** 1D free energy profiles of (A) min($d_{CC}$) and (B) $n_{CC}$ obtained from 2D free energy surfaces shown in Figure S9 by numerically integrating out the other degree of freedom.

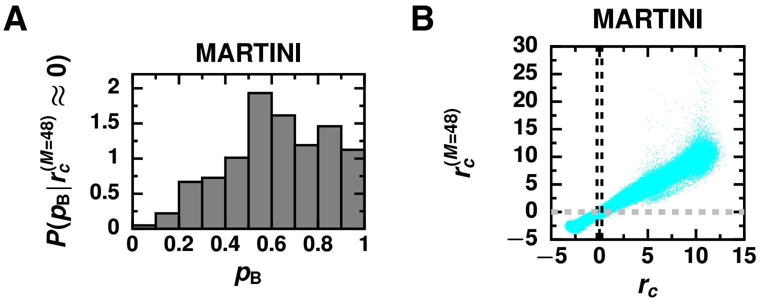


**Figure S11. Reaction coordinate, $r_c^{(M=48)}$, composed of $M = 48$ order parameters, excluding $\min(d_{CC})$ and $n_{CC}$, with coefficients determined using a maximum likelihood approach (Table S4) accurately distinguishes transition states.** (A) Histogram of committor values, $p_B$, for MARTINI configurations found along transition paths with $-0.25 \leq r_c^{(M=48)} \leq 0.25$. (B) Scatter plot of the reaction coordinate, $r_c = \alpha_1 \min(d_{CC}) + \alpha_2 n_{CC} + \alpha_0$ with coefficients determined using the maximum likelihood approach (Table S5) versus $r_c^{(M=48)}$ constructed from $M = 48$ other order parameters for all MARTINI configurations found along transition paths. The black dashed lines outline the values of $r_c$ characteristic of transition states. The gray dashed lines outline the values of $r_c^{(M=48)}$ characteristic of transition states.

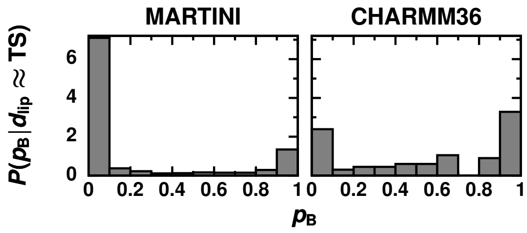


**Figure S12. The COM displacement is not the reaction coordinate for lipid exchange.** Histograms of committor values, $p_B$, for configurations found along transition paths and that have values of $d_{lip}$ typical of transition states (TS). Specifically, MARTINI configurations with $14 \leq d_{lip} \leq 19$ Å and CHARMM36 configurations with $10 \leq d_{lip} \leq 19$ Å are included.

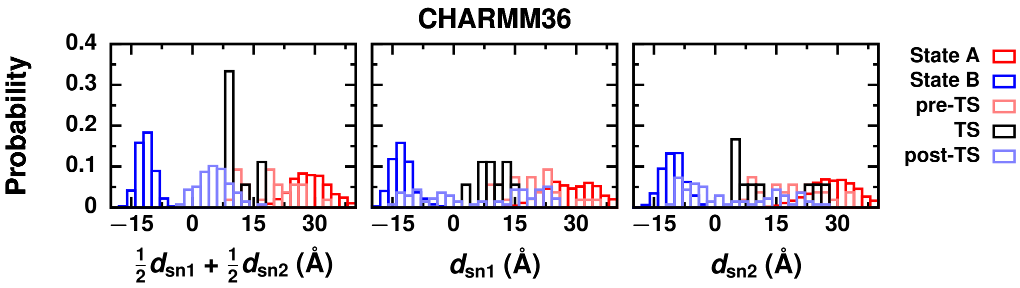


**Figure S13. Displacement-based order parameters do not reliably identify transition states found along transition paths during CHARMM36 simulations when the transition paths are not separated based on pathway.** Probability distributions of $\frac{1}{2}d_{sn1} + \frac{1}{2}d_{sn2}$, $d_{sn1}$, and $d_{sn2}$ from transition paths obtained from CHARMM36 simulations. Distributions are plotted for state A, for state B, and for three ensembles of configurations drawn from transition paths: pre-transition state (pre-TS) configurations identified by $p_B = 0$, transition states (TS) identified by $0.45 \leq p_B \leq 0.55$ for MARTINI and by $0.4 \leq p_B \leq 0.6$ for CHARMM36, and post-transition state (post-TS) configurations identified by $p_B = 1$.

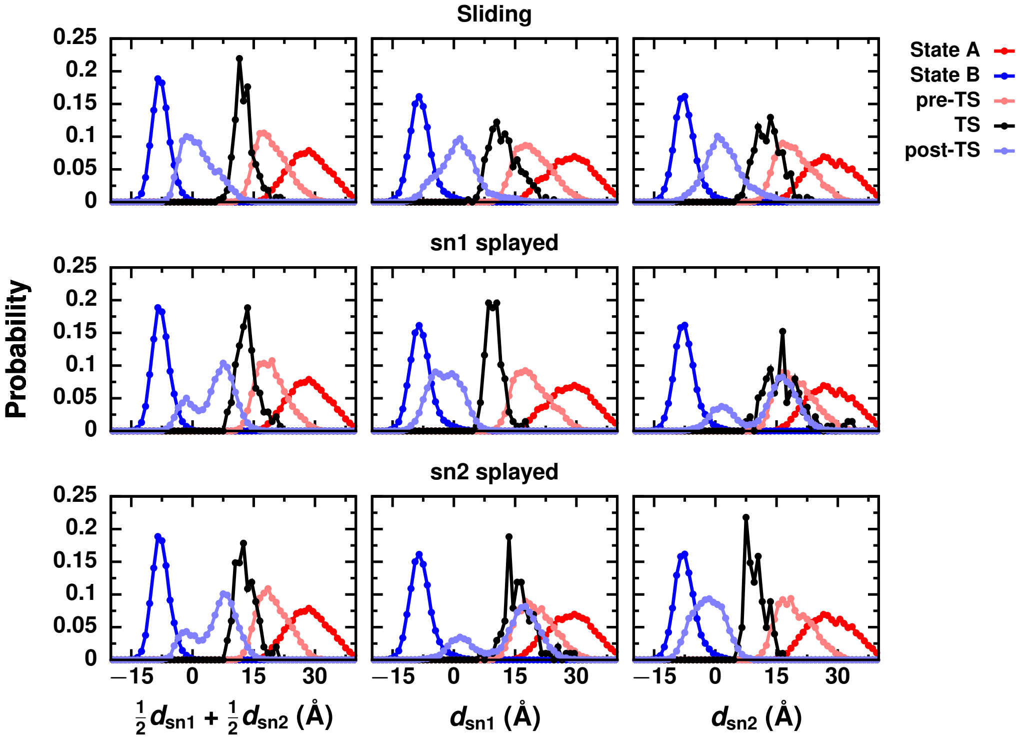


**Figure S14. Displacement-based order parameters focused on the two lipid tails may be pathway specific reaction coordinates.** Probability distributions of $\frac{1}{2}d_{\text{sn1}}+\frac{1}{2}d_{\text{sn2}}$, $d_{\text{sn1}}$, and $d_{\text{sn2}}$ from MARTINI simulations that follow each lipid insertion pathway. Distributions are plotted for state A, for state B, and for three ensembles of configurations drawn from transition paths: pre-transition state (pre-TS) configurations identified by $p_{\text{B}} = 0$, transition states (TS) identified by $0.45 \leq p_{\text{B}} \leq 0.55$ for MARTINI and by $0.4 \leq p_{\text{B}} \leq 0.6$ for CHARMM36, and post-transition state (post-TS) configurations identified by $p_{\text{B}} = 1$.

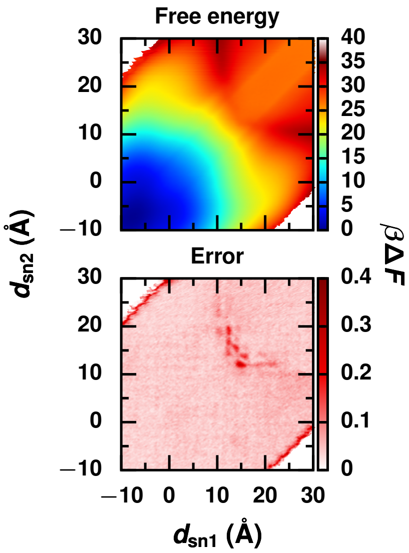


**Figure S15. Error in the free energy surface as a function of the two lipid tail distances is comparable to the barrier height.** (Top panel) Free energy surface as a function of $d_{sn1}$ and $d_{sn2}$ obtained from umbrella sampling simulations using the MARTINI force field. (Bottom panel) Block standard error of the free energy surface calculated from five 200 ns blocks.

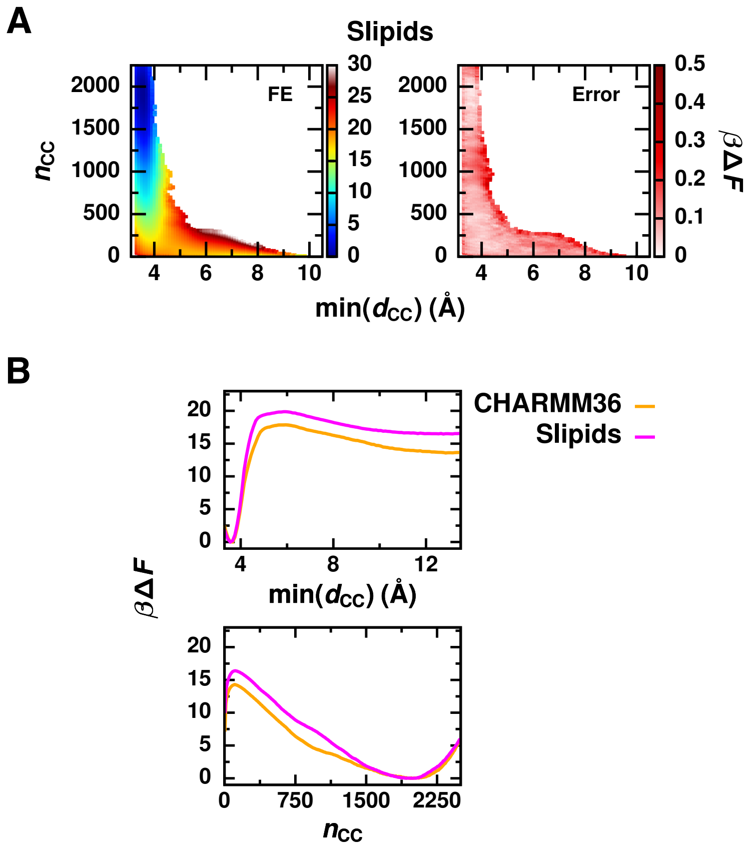


**Figure S16. The height of the free energy barrier for hydrophobic-contact formation/breakage for the Slipids force field is similar but slighly higher than the barrier height for CHARMM36.** (A) (Left) Free energy surfaces as a function of min($d_{CC}$) and $n_{CC}$ obtained from umbrella sampling simulations using the Slipids force field. (Right) Block standard error of the free energy surface calculated from five simulation blocks. (B) 1D free energy profiles of min($d_{CC}$) and $n_{CC}$ for Slipids compared to CHARMM36.

### SUPPLEMENTARY TABLES

**Table S1. Harmonic biases used in umbrella sampling simulations biasing $\min(d_{\mathrm{CC}})$ and $n_{\mathrm{CC}}$ using the MARTINI force field.** Each bias was centered at $(\min(d_{\mathrm{CC}}), n_{\mathrm{CC}}) = (c_0\ \text{Å}, c_1)$. Force constants $k_0$ were used for the harmonic biases on $\min(d_{\mathrm{CC}})$ and are reported in kJ/mol/nm$^2$. Force constants $k_1$ were used for the harmonic biases on $n_{\mathrm{CC}}$ and are reported in kJ/mol/contact$^2$.

| $c_0$ | $c_1$ | $k_0$ | $k_1$ | $c_0$ | $c_1$ | $k_0$ | $k_1$ | $c_0$ | $c_1$ | $k_0$ | $k_1$ | $c_0$ | $c_1$ | $k_0$ | $k_1$ |
|---|---|---|---|---|---|---|---|---|---|---|---|---|---|---|---|
| 0.45 | 0.000 | 1000 | 0.075 | 0.45 | 10.484 | 1000 | 0.075 | 0.45 | 20.968 | 1000 | 0.075 | 0.45 | 31.452 | 1000 | 0.075 |
| 0.45 | 41.935 | 1000 | 0.075 | 0.45 | 52.419 | 1000 | 0.075 | 0.45 | 62.903 | 1000 | 0.075 | 0.45 | 73.387 | 1000 | 0.075 |
| 0.45 | 83.871 | 1000 | 0.075 | 0.45 | 94.355 | 1000 | 0.075 | 0.45 | 104.839 | 1000 | 0.075 | 0.45 | 115.323 | 1000 | 0.075 |
| 0.45 | 125.806 | 1000 | 0.075 | 0.45 | 136.290 | 1000 | 0.075 | 0.45 | 146.774 | 1000 | 0.075 | 0.45 | 157.258 | 1000 | 0.075 |
| 0.45 | 167.742 | 1000 | 0.075 | 0.45 | 178.226 | 1000 | 0.075 | 0.45 | 188.710 | 1000 | 0.075 | 0.45 | 199.194 | 1000 | 0.075 |
| 0.50 | 0.000 | 1000 | 0.075 | 0.50 | 10.484 | 1000 | 0.075 | 0.50 | 20.968 | 1000 | 0.075 | 0.50 | 31.452 | 1000 | 0.075 |
| 0.50 | 41.935 | 1000 | 0.075 | 0.50 | 52.419 | 1000 | 0.075 | 0.50 | 62.903 | 1000 | 0.075 | 0.50 | 73.387 | 1000 | 0.075 |
| 0.50 | 83.871 | 1000 | 0.075 | 0.50 | 94.355 | 1000 | 0.075 | 0.50 | 104.839 | 1000 | 0.075 | 0.50 | 115.323 | 1000 | 0.075 |
| 0.50 | 125.806 | 1000 | 0.075 | 0.50 | 136.290 | 1000 | 0.075 | 0.50 | 146.774 | 1000 | 0.075 | 0.50 | 157.258 | 1000 | 0.075 |
| 0.50 | 167.742 | 1000 | 0.075 | 0.50 | 178.226 | 1000 | 0.075 | 0.50 | 188.710 | 1000 | 0.075 | 0.50 | 199.194 | 1000 | 0.075 |
| 0.55 | 0.000 | 1000 | 0.075 | 0.55 | 10.484 | 1000 | 0.075 | 0.55 | 20.968 | 1000 | 0.075 | 0.55 | 31.452 | 1000 | 0.075 |
| 0.55 | 41.935 | 1000 | 0.075 | 0.55 | 52.419 | 1000 | 0.075 | 0.55 | 62.903 | 1000 | 0.075 | 0.55 | 73.387 | 1000 | 0.075 |
| 0.55 | 83.871 | 1000 | 0.075 | 0.55 | 94.355 | 1000 | 0.075 | 0.55 | 104.839 | 1000 | 0.075 | 0.55 | 115.323 | 1000 | 0.075 |
| 0.55 | 125.806 | 1000 | 0.075 | 0.55 | 136.290 | 1000 | 0.075 | 0.55 | 146.774 | 1000 | 0.075 | 0.55 | 157.258 | 1000 | 0.075 |
| 0.55 | 167.742 | 1000 | 0.075 | 0.55 | 178.226 | 1000 | 0.075 | 0.55 | 188.710 | 1000 | 0.075 | 0.55 | 199.194 | 1000 | 0.075 |
| 0.60 | 0.000 | 1000 | 0.075 | 0.60 | 10.484 | 1000 | 0.075 | 0.60 | 20.968 | 1000 | 0.075 | 0.60 | 31.452 | 1000 | 0.075 |
| 0.60 | 41.935 | 1000 | 0.075 | 0.60 | 52.419 | 1000 | 0.075 | 0.60 | 62.903 | 1000 | 0.075 | 0.60 | 73.387 | 1000 | 0.075 |
| 0.60 | 83.871 | 1000 | 0.075 | 0.60 | 94.355 | 1000 | 0.075 | 0.60 | 104.839 | 1000 | 0.075 | 0.60 | 115.323 | 1000 | 0.075 |
| 0.60 | 125.806 | 1000 | 0.075 | 0.60 | 136.290 | 1000 | 0.075 | 0.60 | 146.774 | 1000 | 0.075 | 0.60 | 157.258 | 1000 | 0.075 |
| 0.60 | 167.742 | 1000 | 0.075 | 0.60 | 178.226 | 1000 | 0.075 | 0.60 | 188.710 | 1000 | 0.075 | 0.60 | 199.194 | 1000 | 0.075 |
| 0.65 | 0.000 | 1000 | 0.075 | 0.65 | 10.484 | 1000 | 0.075 | 0.65 | 20.968 | 1000 | 0.075 | 0.65 | 31.452 | 1000 | 0.075 |
| 0.65 | 41.935 | 1000 | 0.075 | 0.65 | 52.419 | 1000 | 0.075 | 0.65 | 62.903 | 1000 | 0.075 | 0.65 | 73.387 | 1000 | 0.075 |
| 0.65 | 83.871 | 1000 | 0.075 | 0.65 | 94.355 | 1000 | 0.075 | 0.65 | 104.839 | 1000 | 0.075 | 0.65 | 115.323 | 1000 | 0.075 |
| 0.65 | 125.806 | 1000 | 0.075 | 0.65 | 136.290 | 1000 | 0.075 | 0.65 | 146.774 | 1000 | 0.075 | 0.65 | 157.258 | 1000 | 0.075 |
| 0.65 | 167.742 | 1000 | 0.075 | 0.65 | 178.226 | 1000 | 0.075 | 0.65 | 188.710 | 1000 | 0.075 | 0.65 | 199.194 | 1000 | 0.075 |
| 0.70 | 0.000 | 1000 | 0.075 | 0.70 | 10.484 | 1000 | 0.075 | 0.70 | 20.968 | 1000 | 0.075 | 0.70 | 31.452 | 1000 | 0.075 |
| 0.70 | 41.935 | 1000 | 0.075 | 0.70 | 52.419 | 1000 | 0.075 | 0.70 | 62.903 | 1000 | 0.075 | 0.70 | 73.387 | 1000 | 0.075 |
| 0.70 | 83.871 | 1000 | 0.075 | 0.70 | 94.355 | 1000 | 0.075 | 0.70 | 104.839 | 1000 | 0.075 | 0.70 | 115.323 | 1000 | 0.075 |
| 0.70 | 125.806 | 1000 | 0.075 | 0.70 | 136.290 | 1000 | 0.075 | 0.70 | 146.774 | 1000 | 0.075 | 0.70 | 157.258 | 1000 | 0.075 |
| 0.70 | 167.742 | 1000 | 0.075 | 0.70 | 178.226 | 1000 | 0.075 | 0.70 | 188.710 | 1000 | 0.075 | 0.70 | 199.194 | 1000 | 0.075 |
| 0.75 | 0.000 | 1000 | 0.075 | 0.75 | 10.484 | 1000 | 0.075 | 0.75 | 20.968 | 1000 | 0.075 | 0.75 | 31.452 | 1000 | 0.075 |
| 0.75 | 41.935 | 1000 | 0.075 | 0.75 | 52.419 | 1000 | 0.075 | 0.80 | 0.000 | 1000 | 0.075 | 0.80 | 10.484 | 1000 | 0.075 |
| 0.80 | 20.968 | 1000 | 0.075 | 0.80 | 31.452 | 1000 | 0.075 | 0.80 | 41.935 | 1000 | 0.075 | 0.80 | 52.419 | 1000 | 0.075 |
| 0.85 | 0.000 | 1000 | 0.075 | 0.85 | 10.484 | 1000 | 0.075 | 0.85 | 20.968 | 1000 | 0.075 | 0.85 | 31.452 | 1000 | 0.075 |
| 0.85 | 41.935 | 1000 | 0.075 | 0.85 | 52.419 | 1000 | 0.075 | 0.90 | 0.000 | 1000 | 0.075 | 0.90 | 10.484 | 1000 | 0.075 |
| 0.90 | 20.968 | 1000 | 0.075 | 0.90 | 31.452 | 1000 | 0.075 | 0.90 | 41.935 | 1000 | 0.075 | 0.90 | 52.419 | 1000 | 0.075 |
| 0.95 | 0.000 | 1000 | 0.075 | 0.95 | 10.484 | 1000 | 0.075 | 0.95 | 20.968 | 1000 | 0.075 | 0.95 | 31.452 | 1000 | 0.075 |
| 0.95 | 41.935 | 1000 | 0.075 | 0.95 | 52.419 | 1000 | 0.075 | 1.00 | 0.000 | 1000 | 0.075 | 1.00 | 10.484 | 1000 | 0.075 |
| 1.00 | 20.968 | 1000 | 0.075 | 1.00 | 31.452 | 1000 | 0.075 | 1.00 | 41.935 | 1000 | 0.075 | 1.00 | 52.419 | 1000 | 0.075 |
| 1.05 | 0.000 | 1000 | 0.075 | 1.05 | 10.484 | 1000 | 0.075 | 1.05 | 20.968 | 1000 | 0.075 | 1.05 | 31.452 | 1000 | 0.075 |
| 1.05 | 41.935 | 1000 | 0.075 | 1.05 | 52.419 | 1000 | 0.075 | 1.10 | 0.000 | 1000 | 0.075 | 1.10 | 10.484 | 1000 | 0.075 |
| 1.10 | 20.968 | 1000 | 0.075 | 1.10 | 31.452 | 1000 | 0.075 | 1.10 | 41.935 | 1000 | 0.075 | 1.10 | 52.419 | 1000 | 0.075 |
| 1.15 | 0.000 | 1000 | 0.075 | 1.15 | 10.484 | 1000 | 0.075 | 1.15 | 20.968 | 1000 | 0.075 | 1.15 | 31.452 | 1000 | 0.075 |
| 1.15 | 41.935 | 1000 | 0.075 | 1.15 | 52.419 | 1000 | 0.075 | 1.20 | 0.000 | 1000 | 0.075 | 1.20 | 10.484 | 1000 | 0.075 |
| 1.20 | 20.968 | 1000 | 0.075 | 1.20 | 31.452 | 1000 | 0.075 | 1.20 | 41.935 | 1000 | 0.075 | 1.20 | 52.419 | 1000 | 0.075 |
| 1.25 | 0.000 | 1000 | 0.000 | 1.30 | 0.000 | 1000 | 0.000 | 1.35 | 0.000 | 1000 | 0.000 | 1.40 | 0.000 | 1000 | 0.000 |
| 1.45 | 0.000 | 1000 | 0.000 | 1.50 | 0.000 | 1000 | 0.000 | 1.55 | 0.000 | 1000 | 0.000 | 1.60 | 0.000 | 1000 | 0.000 |
| 1.65 | 0.000 | 1000 | 0.000 | 1.70 | 0.000 | 1000 | 0.000 | 1.75 | 0.000 | 1000 | 0.000 | 1.80 | 0.000 | 1000 | 0.000 |
| 1.85 | 0.000 | 1000 | 0.000 | 1.90 | 0.000 | 1000 | 0.000 | 1.95 | 0.000 | 1000 | 0.000 | 2.00 | 0.000 | 1000 | 0.000 |
| 0.47 | 209.677 | 0 | 0.075 | 0.47 | 220.161 | 0 | 0.075 | 0.47 | 230.645 | 0 | 0.075 | 0.47 | 241.129 | 0 | 0.075 |
| 0.47 | 251.613 | 0 | 0.075 | 0.47 | 262.097 | 0 | 0.075 | 0.47 | 272.581 | 0 | 0.075 | 0.47 | 283.065 | 0 | 0.075 |
| 0.47 | 293.548 | 0 | 0.075 | 0.47 | 304.032 | 0 | 0.075 | 0.47 | 314.516 | 0 | 0.075 | 0.47 | 325.000 | 0 | 0.075 |

**Table S2. Harmonic biases used in umbrella sampling simulations biasing min($d_{\mathrm{CC}}$) and $n_{\mathrm{CC}}$ using the CHARMM36 and Slipids force fields.** Each bias was centered at $(\min(d_{\mathrm{CC}}), n_{\mathrm{CC}}) = (c_0\ \text{Å}, c_1)$. Force constants $k_0$ were used for the harmonic biases on min($d_{\mathrm{CC}}$) and are reported in kJ/mol/nm$^2$. Force constants $k_1$ were used for the harmonic biases on $n_{\mathrm{CC}}$ and are reported in kJ/mol/contact$^2$.

| $c_0$ | $c_1$ | $k_0$ | $k_1$ | $c_0$ | $c_1$ | $k_0$ | $k_1$ | $c_0$ | $c_1$ | $k_0$ | $k_1$ | $c_0$ | $c_1$ | $k_0$ | $k_1$ |
|---|---|---|---|---|---|---|---|---|---|---|---|---|---|---|---|
| 0.32 | 0.000 | 1000 | 0.002 | 0.32 | 64.516 | 1000 | 0.002 | 0.32 | 129.032 | 1000 | 0.002 | 0.32 | 193.548 | 1000 | 0.002 |
| 0.32 | 258.065 | 1000 | 0.002 | 0.32 | 322.581 | 1000 | 0.002 | 0.32 | 387.097 | 1000 | 0.002 | 0.32 | 451.613 | 1000 | 0.002 |
| 0.32 | 516.129 | 1000 | 0.002 | 0.32 | 580.645 | 1000 | 0.002 | 0.32 | 645.161 | 1000 | 0.002 | 0.32 | 709.677 | 1000 | 0.002 |
| 0.32 | 774.194 | 1000 | 0.002 | 0.32 | 838.710 | 1000 | 0.002 | 0.32 | 903.226 | 1000 | 0.002 | 0.32 | 967.742 | 1000 | 0.002 |
| 0.32 | 1032.258 | 1000 | 0.002 | 0.32 | 1096.774 | 1000 | 0.002 | 0.32 | 1161.290 | 1000 | 0.002 | 0.32 | 1225.806 | 1000 | 0.002 |
| 0.32 | 1290.323 | 1000 | 0.002 | 0.37 | 0.000 | 1000 | 0.002 | 0.37 | 64.516 | 1000 | 0.002 | 0.37 | 129.032 | 1000 | 0.002 |
| 0.37 | 193.548 | 1000 | 0.002 | 0.37 | 258.065 | 1000 | 0.002 | 0.37 | 322.581 | 1000 | 0.002 | 0.37 | 387.097 | 1000 | 0.002 |
| 0.37 | 451.613 | 1000 | 0.002 | 0.37 | 516.129 | 1000 | 0.002 | 0.37 | 580.645 | 1000 | 0.002 | 0.37 | 645.161 | 1000 | 0.002 |
| 0.37 | 709.677 | 1000 | 0.002 | 0.37 | 774.194 | 1000 | 0.002 | 0.37 | 838.710 | 1000 | 0.002 | 0.37 | 903.226 | 1000 | 0.002 |
| 0.37 | 967.742 | 1000 | 0.002 | 0.37 | 1032.258 | 1000 | 0.002 | 0.37 | 1096.774 | 1000 | 0.002 | 0.37 | 1161.290 | 1000 | 0.002 |
| 0.37 | 1225.806 | 1000 | 0.002 | 0.37 | 1290.323 | 1000 | 0.002 | 0.42 | 0.000 | 1000 | 0.002 | 0.42 | 64.516 | 1000 | 0.002 |
| 0.42 | 129.032 | 1000 | 0.002 | 0.42 | 193.548 | 1000 | 0.002 | 0.42 | 258.065 | 1000 | 0.002 | 0.42 | 322.581 | 1000 | 0.002 |
| 0.42 | 387.097 | 1000 | 0.002 | 0.42 | 451.613 | 1000 | 0.002 | 0.42 | 516.129 | 1000 | 0.002 | 0.42 | 580.645 | 1000 | 0.002 |
| 0.42 | 645.161 | 1000 | 0.002 | 0.42 | 709.677 | 1000 | 0.002 | 0.42 | 774.194 | 1000 | 0.002 | 0.42 | 838.710 | 1000 | 0.002 |
| 0.42 | 903.226 | 1000 | 0.002 | 0.42 | 967.742 | 1000 | 0.002 | 0.42 | 1032.258 | 1000 | 0.002 | 0.42 | 1096.774 | 1000 | 0.002 |
| 0.42 | 1161.290 | 1000 | 0.002 | 0.42 | 1225.806 | 1000 | 0.002 | 0.42 | 1290.323 | 1000 | 0.002 | 0.47 | 0.000 | 1000 | 0.002 |
| 0.47 | 64.516 | 1000 | 0.002 | 0.47 | 129.032 | 1000 | 0.002 | 0.47 | 193.548 | 1000 | 0.002 | 0.47 | 258.065 | 1000 | 0.002 |
| 0.47 | 322.581 | 1000 | 0.002 | 0.47 | 387.097 | 1000 | 0.002 | 0.47 | 451.613 | 1000 | 0.002 | 0.47 | 516.129 | 1000 | 0.002 |
| 0.47 | 580.645 | 1000 | 0.002 | 0.47 | 645.161 | 1000 | 0.002 | 0.47 | 709.677 | 1000 | 0.002 | 0.47 | 774.194 | 1000 | 0.002 |
| 0.47 | 838.710 | 1000 | 0.002 | 0.47 | 903.226 | 1000 | 0.002 | 0.47 | 967.742 | 1000 | 0.002 | 0.47 | 1032.258 | 1000 | 0.002 |
| 0.47 | 1096.774 | 1000 | 0.002 | 0.47 | 1161.290 | 1000 | 0.002 | 0.47 | 1225.806 | 1000 | 0.002 | 0.47 | 1290.323 | 1000 | 0.002 |
| 0.52 | 0.000 | 1000 | 0.002 | 0.52 | 64.516 | 1000 | 0.002 | 0.52 | 129.032 | 1000 | 0.002 | 0.52 | 193.548 | 1000 | 0.002 |
| 0.52 | 258.065 | 1000 | 0.002 | 0.52 | 322.581 | 1000 | 0.002 | 0.52 | 387.097 | 1000 | 0.002 | 0.52 | 451.613 | 1000 | 0.002 |
| 0.52 | 516.129 | 1000 | 0.002 | 0.52 | 580.645 | 1000 | 0.002 | 0.52 | 645.161 | 1000 | 0.002 | 0.52 | 709.677 | 1000 | 0.002 |
| 0.52 | 774.194 | 1000 | 0.002 | 0.52 | 838.710 | 1000 | 0.002 | 0.52 | 903.226 | 1000 | 0.002 | 0.52 | 967.742 | 1000 | 0.002 |
| 0.52 | 1032.258 | 1000 | 0.002 | 0.52 | 1096.774 | 1000 | 0.002 | 0.52 | 1161.290 | 1000 | 0.002 | 0.52 | 1225.806 | 1000 | 0.002 |
| 0.52 | 1290.323 | 1000 | 0.002 | 0.57 | 0.000 | 1000 | 0.002 | 0.57 | 64.516 | 1000 | 0.002 | 0.57 | 129.032 | 1000 | 0.002 |
| 0.57 | 193.548 | 1000 | 0.002 | 0.57 | 258.065 | 1000 | 0.002 | 0.57 | 322.581 | 1000 | 0.002 | 0.57 | 387.097 | 1000 | 0.002 |
| 0.57 | 451.613 | 1000 | 0.002 | 0.57 | 516.129 | 1000 | 0.002 | 0.57 | 580.645 | 1000 | 0.002 | 0.57 | 645.161 | 1000 | 0.002 |
| 0.57 | 709.677 | 1000 | 0.002 | 0.57 | 774.194 | 1000 | 0.002 | 0.57 | 838.710 | 1000 | 0.002 | 0.57 | 903.226 | 1000 | 0.002 |
| 0.57 | 967.742 | 1000 | 0.002 | 0.57 | 1032.258 | 1000 | 0.002 | 0.57 | 1096.774 | 1000 | 0.002 | 0.57 | 1161.290 | 1000 | 0.002 |
| 0.57 | 1225.806 | 1000 | 0.002 | 0.57 | 1290.323 | 1000 | 0.002 | 0.62 | 0.000 | 1000 | 0.002 | 0.62 | 64.516 | 1000 | 0.002 |
| 0.62 | 129.032 | 1000 | 0.002 | 0.62 | 193.548 | 1000 | 0.002 | 0.62 | 258.065 | 1000 | 0.002 | 0.67 | 0.000 | 1000 | 0.002 |
| 0.67 | 64.516 | 1000 | 0.002 | 0.67 | 129.032 | 1000 | 0.002 | 0.67 | 193.548 | 1000 | 0.002 | 0.67 | 258.065 | 1000 | 0.002 |
| 0.72 | 0.000 | 1000 | 0.002 | 0.72 | 64.516 | 1000 | 0.002 | 0.72 | 129.032 | 1000 | 0.002 | 0.72 | 193.548 | 1000 | 0.002 |
| 0.72 | 258.065 | 1000 | 0.002 | 0.77 | 0.000 | 1000 | 0.002 | 0.77 | 64.516 | 1000 | 0.002 | 0.77 | 129.032 | 1000 | 0.002 |
| 0.77 | 193.548 | 1000 | 0.002 | 0.77 | 258.065 | 1000 | 0.002 | 0.82 | 0.000 | 1000 | 0.002 | 0.82 | 64.516 | 1000 | 0.002 |
| 0.82 | 129.032 | 1000 | 0.002 | 0.82 | 193.548 | 1000 | 0.002 | 0.82 | 258.065 | 1000 | 0.002 | 0.87 | 0.000 | 1000 | 0.000 |
| 0.92 | 0.000 | 1000 | 0.000 | 0.97 | 0.000 | 1000 | 0.000 | 1.02 | 0.000 | 1000 | 0.000 | 1.07 | 0.000 | 1000 | 0.000 |
| 1.12 | 0.000 | 1000 | 0.000 | 1.17 | 0.000 | 1000 | 0.000 | 1.22 | 0.000 | 1000 | 0.000 | 1.27 | 0.000 | 1000 | 0.000 |
| 1.32 | 0.000 | 1000 | 0.000 | 1.37 | 0.000 | 1000 | 0.000 | 1.42 | 0.000 | 1000 | 0.000 | 1.47 | 0.000 | 1000 | 0.000 |
| 1.52 | 0.000 | 1000 | 0.000 | 1.57 | 0.000 | 1000 | 0.000 | 1.62 | 0.000 | 1000 | 0.000 | 1.67 | 0.000 | 1000 | 0.000 |
| 1.72 | 0.000 | 1000 | 0.000 | 1.77 | 0.000 | 1000 | 0.000 | 1.82 | 0.000 | 1000 | 0.000 | 1.87 | 0.000 | 1000 | 0.000 |
| 0.37 | 1354.839 | 0 | 0.002 | 0.37 | 1419.355 | 0 | 0.002 | 0.37 | 1483.871 | 0 | 0.002 | 0.37 | 1548.387 | 0 | 0.002 |
| 0.37 | 1612.903 | 0 | 0.002 | 0.37 | 1677.419 | 0 | 0.002 | 0.37 | 1741.935 | 0 | 0.002 | 0.37 | 1806.452 | 0 | 0.002 |
| 0.37 | 1870.968 | 0 | 0.002 | 0.37 | 1935.484 | 0 | 0.002 | 0.37 | 2000.000 | 0 | 0.002 | 0.37 | 2064.516 | 0 | 0.002 |
| 0.37 | 2129.032 | 0 | 0.002 | 0.37 | 2193.548 | 0 | 0.002 | 0.37 | 2258.064 | 0 | 0.002 | 0.37 | 2322.580 | 0 | 0.002 |
| 0.37 | 2387.096 | 0 | 0.002 | 0.37 | 2451.612 | 0 | 0.002 | 0.37 | 2516.128 | 0 | 0.002 | 0.42 | 1354.839 | 1000 | 0.002 |
| 0.42 | 1419.355 | 1000 | 0.002 | 0.42 | 1483.871 | 1000 | 0.002 | 0.42 | 1548.387 | 1000 | 0.002 | 0.42 | 1612.903 | 1000 | 0.002 |
| 0.42 | 1677.419 | 1000 | 0.002 | 0.42 | 1741.935 | 1000 | 0.002 | 0.42 | 1806.452 | 1000 | 0.002 | 0.42 | 1870.968 | 1000 | 0.002 |
| 0.42 | 1935.484 | 1000 | 0.002 | 0.42 | 2000.000 | 1000 | 0.002 | | | | | | | | |

**Table S3. Frequency of each insertion pathway.**

| Pathway | MARTINI | CHARMM36 |
|---|---|---|
| Sliding | 51% | 20% |
| sn1 splayed | 27% | 40% |
| sn2 splayed | 22% | 40% |

**Table S4. Construction of a reaction coordinate, $r_c^{(M=48)}$, excluding $\min(d_{\mathrm{CC}})$ and $n_{\mathrm{CC}}$ using a maximum likelihood approach.** The best reaction coordinates for the MARTINI model with $M = 1, 2$, and 48 are reported. To determine the best reaction coordinate with $M = 2$, only linear combinations with $d_{\mathrm{lip}}$, $d_{\mathrm{sn1}}$, or $d_{\mathrm{sn2}}$ as one of the order parameters were considered. To determine the best reaction coordinate with $M = 3$, only linear combinations with either (1) $d_{\mathrm{lip}}$ and $\rho^{(\mathrm{cyl})}_{\mathrm{philic,lip}}$ or (2) $d_{\mathrm{sn1}}$ and $d_{\mathrm{sn2}}$ were considered. All reported reaction coordinates are constructed from order parameters scaled to the range [0,1]. Scaled order parameters are denoted with a tilde to distinguish them from the unscaled values.

| $M$ | $r_c^{(M=48)}$ | $\ln \mathcal{L}$ |
|---|---|---|
| 1 | $-12.635\tilde{d}_{\mathrm{lip}} + 8.060$ | −937984 |
| 2 | $-13.583\tilde{d}_{\mathrm{lip}} - 5.663\tilde{\rho}^{(\mathrm{cyl})}_{\mathrm{philic,lip}} + 11.296$ | −681129 |
| 3 | $-11.889\tilde{d}_{\mathrm{lip}} - 4.411\tilde{\rho}^{(\mathrm{cyl})}_{\mathrm{philic,lip}} + 18.892\tilde{\rho}^{(\mathrm{cyl})}_{\mathrm{phobic,sn1}} + 7.680$ | −596942 |
| 48 | $-3.919\tilde{d}_{\mathrm{lip}} - 2.455\tilde{d}_{\mathrm{sn1}} - 1.529\tilde{d}_{\mathrm{sn2}} - 0.116\tilde{n}^{(\mathrm{lip})}_{\mathrm{w,NC3}} - 0.054\tilde{n}^{(\mathrm{lip})}_{\mathrm{w,PO4}} - 0.363\tilde{n}^{(\mathrm{lip})}_{\mathrm{w,GL1}} - 0.436\tilde{n}^{(\mathrm{lip})}_{\mathrm{w,GL2}} + 0.051\tilde{n}^{(\mathrm{lip})}_{\mathrm{w,C1A}} + 0.278\tilde{n}^{(\mathrm{lip})}_{\mathrm{w,C2A}} + 0.231\tilde{n}^{(\mathrm{lip})}_{\mathrm{w,C3A}} - 0.028\tilde{n}^{(\mathrm{lip})}_{\mathrm{w,C1B}} + 0.208\tilde{n}^{(\mathrm{lip})}_{\mathrm{w,C2B}} + 0.099\tilde{n}_{\mathrm{w,C3B}} + 0.174\tilde{n}^{(\mathrm{mem})}_{\mathrm{w,NC3}} - 0.256\tilde{n}^{(\mathrm{mem})}_{\mathrm{w,PO4}} - 0.90\tilde{n}^{(\mathrm{mem})}_{\mathrm{w,GL1}} - 0.401\tilde{n}^{(\mathrm{mem})}_{\mathrm{w,GL2}} - 0.001\tilde{n}^{(\mathrm{mem})}_{\mathrm{w,C1A}} + 0.188\tilde{n}^{(\mathrm{mem})}_{\mathrm{w,C2A}} + 0.007\tilde{n}^{(\mathrm{mem})}_{\mathrm{w,C3A}} - 0.023\tilde{n}^{(\mathrm{mem})}_{\mathrm{w,C1B}} + 0.043\tilde{n}^{(\mathrm{mem})}_{\mathrm{w,C2B}} + 0.048\tilde{n}^{(\mathrm{mem})}_{\mathrm{w,C3B}} + 1.447\langle\delta\tilde{z}_{\mathrm{phos}}\rangle_{\mathrm{mem}} + 0.813\langle\tilde{n}_{\mathrm{ET}}\rangle_{\mathrm{mem}} + 0.169\tilde{A}_{\mathrm{defect,sn1}} - 0.013\tilde{A}_{\mathrm{defect,sn2}} + 0.359\min(\tilde{d}_{\mathrm{HT,sn1}}) + 0.255\min(\tilde{d}_{\mathrm{HT,sn2}}) - 0.572\tilde{\rho}^{\mathrm{cyl})}_{\mathrm{w,lip}} - 5.839\tilde{\rho}^{(\mathrm{cyl})}_{\mathrm{w,sn1}} - 5.215\tilde{\rho}^{(\mathrm{cyl})}_{\mathrm{w,sn2}} - 0.521\tilde{\rho}^{(\mathrm{cyl})}_{\mathrm{lip,lip}} - 0.865\tilde{\rho}^{(\mathrm{cyl})}_{\mathrm{philic,lip}} + 1.499\tilde{\rho}^{(\mathrm{cyl})}_{\mathrm{phobic,lip}} + 4.337\tilde{\rho}^{(\mathrm{cyl})}_{\mathrm{lip,sn1}} + 2.885\tilde{\rho}^{(\mathrm{cyl})}_{\mathrm{lip,sn2}} - 1.264\tilde{\rho}^{(\mathrm{cyl})}_{\mathrm{philic,sn1}} - 1.504\tilde{\rho}^{(\mathrm{cyl})}_{\mathrm{philic,sn2}} + 17.591\tilde{\rho}^{(\mathrm{cyl})}_{\mathrm{phobic,sn1}} + 18.334\tilde{\rho}^{(\mathrm{cyl})}_{\mathrm{phobic,sn2}} + 7.904\tilde{n}^{(\mathrm{cyl})}_{\mathrm{w,sn1}} + 7.161\tilde{n}^{(\mathrm{cyl})}_{\mathrm{w,sn2}} - 2.827\tilde{n}^{(\mathrm{cyl})}_{\mathrm{phobic,sn1}} - 4.252\tilde{n}^{(\mathrm{cyl})}_{\mathrm{phobic,sn2}} - 0.442\tilde{n}^{(\mathrm{cyl})}_{\mathrm{all,sn1}} + 0.443\tilde{n}^{(\mathrm{cyl})}_{\mathrm{all,sn2}} - 1.184\tilde{E}_{\mathrm{int}} + 5.291$ | −415253 |

**Table S5. Construction of a reaction coordinate that can include $n_{\mathrm{CC}}$ and $\min(d_{\mathrm{CC}})$ using a maximum likelihood approach demonstrates that a linear combination of $n_{\mathrm{CC}}$ and $\min(d_{\mathrm{CC}})$ is the ideal reaction coordinate.** To construct $r_c$, 54 and 32 order parameters were considered for the MARTINI and CHARMM36 models, respectively. The best reaction coordinates with $M = 1$ and 2 are reported. To determine the best reaction coordinate with $M = 2$, only linear combinations with $\min(d_{\mathrm{CC}})$ or $n_{\mathrm{CC}}$ (and tail specific variants) as one of the order parameters were considered. All reported reaction coordinates are constructed from order parameters scaled to the range [0,1]. Scaled order parameters are denoted with a tilde to distinguish them from the unscaled values.

| $M$ | $r_c$ | $\ln \mathcal{L}$ |
|---|---|---|
| | **MARTINI** | |
| 1 | $24.917\tilde{n}_{\mathrm{CC}} - 1.922$ | −419147 |
| 2 | $-3.136\min(\tilde{d}_{\mathrm{CC}}) + 13.643\tilde{n}_{\mathrm{CC}} - 0.516$ | −399237 |
| | **CHARMM36** | |
| 1 | $18.583\tilde{n}_{\mathrm{CC}} - 1.217$ | −584 |
| 2 | $-3.982\min(\tilde{d}_{\mathrm{CC}}) + 8.102\tilde{n}_{\mathrm{CC}} - 0.151$ | −515 |
